# CNTNAP4 signaling regulates osteosarcoma disease progression

**DOI:** 10.1101/2022.08.26.505419

**Authors:** Qizhi Qin, Sowmya Ramesh, Mario Gomez-Salazar, Lingke Zhong, Masnsen Cherief, Aniket Pratapneni, Carol D. Morris, Edward F. McCarthy, Xinli Zhang, Aaron W. James

## Abstract

Improved treatment strategies for sarcoma rely on clarification of the molecular mediators of disease progression. Recently, we reported that the secreted glycoprotein NELL-1 modulates osteosarcoma (OS) disease progression in part via altering the sarcomatous extracellular matrix (ECM) and cell-ECM interactions. Of known NELL-1 interactor proteins, Contactin-associated protein-like 4 (Cntnap4) encodes a member of the neurexin superfamily of transmembrane molecules best known for its presynaptic functions in the central nervous system. Here, CRISPR/Cas9 gene deletion of *CNTNAP4* reduced OS tumor growth, sarcoma-associated angiogenesis, and pulmonary metastases. *CNTNAP4* knockout (KO) in OS tumor cells largely phenocopied the effects of *NELL-1* KO, including reductions in sarcoma cell attachment, migration, and invasion. Further, *CNTNAP4* KO cells were found to be unresponsive to the effects of NELL-1 treatment. Transcriptomic analysis combined with protein phospho-array demonstrated notable reductions in the MAPK/ERK signaling cascade with *CNTNAP4* deletion, and the ERK1/2 agonist isoproterenol restored cell functions among *CNTNAP4* KO tumor cells. Finally, human primary cells and tissues in combination with sequencing datasets confirmed the significance of *CNTNAP4* signaling in human sarcomas. In summary, our findings demonstrate the biological importance of NELL-1/CNTNAP4 signaling axis in disease progression of human sarcomas and suggest that targeting the NELL-1/CNTNAP4 signaling pathway represents a strategy with potential therapeutic benefit in sarcoma patients.

## Introduction

Osteosarcoma (OS) is the most common primary bone malignancy and post-radiation sarcoma ^1, 2^, and is the most common bone sarcoma in children and adolescents ^1^. Despite advances in therapy, the overall 5-year survival rate remains poor and the prognosis for patients with metastatic disease is grim ^3, 4^. Prior observations identified high expression of the osteoblast-related protein NELL-1 (NEL-like molecule-1) in human OS specimens ^5^. Indeed, we recently observed that the secreted protein NELL-1 regulates osteosarcomagenesis and OS disease progression, associated with alteration in the sarcoma matrisome and FAK/Src signaling activation ^6^. Nevertheless, NELL-1 is known to bind to and activate multiple receptors ^7^, and the ligand-receptor interactions in NELL-1 regulation of OS biology remain unexplored.

Cntnap4 (Contactin-associated protein-like 4, also known as Caspr4) is a member of the neurexin superfamily receptor that is highly enriched in interneurons ^8^. Cntnap4 interacts with presynaptic proteins to mediate neuronal communication and differentiation ^8, 9^. Recently, a ligand/receptor-like interaction between NELL-1 and Cntnap4 was reported, which plays a critical role in NELL-1 cellular response, including FAK (focal adhesion kinase) and canonical Wnt signaling activation in osteoblasts ^7^. *Cntnap4* knockdown replicated the effects of *Nell1* knockdown *in vitro* in osteoblast cell lines, and transgenic lineage restricted *Cntnap4* knockout (KO) animals phenocopied *Nell1* KO mice ^7^. This molecular data has shown that NELL-1 is a high-affinity ligand for CNTNAP4 and implies that autocrine NELL-1/CNTNAP4 signaling within sarcoma cells could be a biologically relevant axis that regulates OS disease progression.

In this study, we sought to define the regulatory role of *CNTNAP4* in OS pathogenesis. Results showed that CRISPR/Cas9 mediated *CNTNAP4* gene deletion markedly reduced the aggressive phenotype across human OS cell lines. *CNTNAP4* KO significantly slowed OS disease progression, blunted metastatic potential, and reduced angiogenesis in a xenograft OS model. Bulk RNA Sequencing (RNA-Seq) and protein phospho-array demonstrated downregulation of key genes in the Mitogen-activated protein kinase (MAPK) signaling pathway in *CNTNAP4* KO cells. These findings in mouse models suggest that CNTNAP4 signaling positively regulates multiple aspects of OS disease progression in part via perturbations in the Ras-MAPK/ERK signaling cascade, and that targeting NELL-1/CNTNAP4 signaling may represent an alternative therapeutic approach.

## Results

### Cell autonomous effects of *CNTNAP4* gene deletion in osteosarcoma

To assess the role of *CNTNAP4* in OS, we first examined *CNTNAP4* gene expression across different human OS cell lines and observed the highest expression of *CNTNAP4* in 143B cells (**Fig. 1a**). Next, *CNTNAP4* KO clones were generated from the 143B OS cell line using CRISPR/Cas9. *CNTNAP4* gene deletion in two clones was confirmed using qRT-PCR, western blot, and the T7 endonuclease I assay (**Fig. 1b, c**, **Supplementary Figure 1, 2**). When compared to vector control (VC), multiple cellular effects were observed among *CNTNAP4* KO 143B clones, including reduced proliferation (**Fig. 1c**, mean 37.3% reduction at 72 h), reduced cell attachment (**Fig. 1d, e**, mean 19.6% reduction at 5 h), reduced cell migration (**Fig. 1f,** **g, 32**.0-36.7% reduction across clones), and reduced cellular invasion (**Fig. 1h, i**, 54.1-60.7% reduction). Similar findings were observed with a polyclonal *CNTNAP4* KO cell preparation either in the 143B or Saos2 OS cell lines (15.8-49.5% reduction in proliferation, 32.2-44.7% reduction in attachment, **Supplementary Figure 3, 4**). Next, we determined whether *CNTNAP4* KO cells showed deficient responsiveness to NELL-1. For this purpose, rNELL-1 protein coating was applied and the attachment rates of control or *CNTNAP4* KO 143B cells was assayed (**Fig. 1j, k**). Results showed an increase in attachment rates among control 143B cells in agreement with prior reports ^10^, which was not seen among *CNTNAP4* KO cells. Thus, *CNTNAP4* plays a crucial role in maintaining cellular proliferation, attachment, migration, and invasion potential in OS cells, at least in part mediated by NELL-1 / CNTNAP4 interaction.

**Figure 1.**
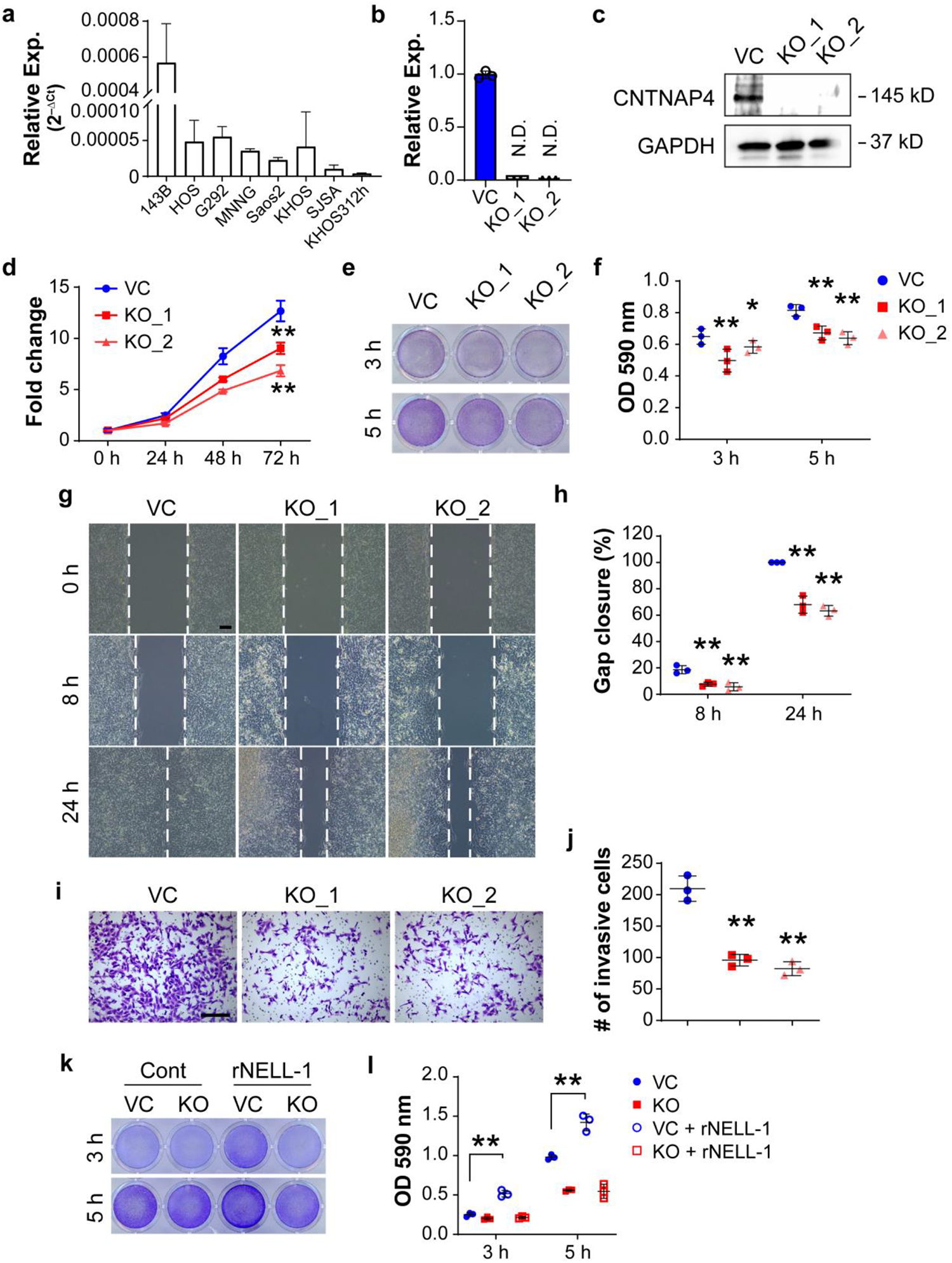
Cell autonomous effects of CRISPR/Cas9-mediated *CNTNAP4* gene deletion in human osteosarcoma cell lines. **(a)** Relative *CNTNAP4* mRNA expression across osteosarcoma cell lines. (**b**) Relative expression of *CNTNAP4* in two clones compared with vector control (VC) by qPCR. **(c)** Western blot of CNTNAP4 in control 143B cells (VC) and knockout clones (KO). **(d)** MTS proliferation assay among *CNTNAP4* KO single cell clones at 24, 48, and 72 h. **(e, f)** Attachment assay at 3 and 5 h, assessed by (**e**) crystal violet staining and (**f**) quantification. (**g, h**) Migration assay at 8 and 24 h assessed by (**g**) scratch wound healing assay and (**h**) quantification. **(i, j)** Transwell invasion assay at 24 h, including (**i**) representative images and (**j**) quantification. (**k, l**) Effects of recombinant NELL1 (5 μg/cm^2^) on 143B OS cells with or without *CNTNAP4* KO. Attachment assay assessed at 3 and 5 h by (**k**) crystal violet staining and (**l**) quantification. Data shown as mean ± 1 SD, with dots representing individual data points. All experiments performed in triplicates, with results from a single replicate shown. ND: Not detected. \**P*<0.05; \*\**P*<0.01 in comparison to VC. Scale bars: 100 µm.

### *CNTNAP4* knockout mitigates OS disease progression

To further assess the effect of *CNTNAP4* gene deletion, we next utilized an orthotopic xenograft model in which *CNTNAP4* KO or control 143B cells were implanted into NOD scid mice (**Fig. 2**). First, serial caliper measurements assessed tumor growth (**Fig. 2a**). A clear reduction of tumor size was observed in *CNTNAP4* KO group when compared to the control group (49.1% reduction at 28 d post-implantation). High resolution XR imaging was performed and confirmed a 48.0% reduction in tumor size at 28 d post-implantation (**Fig. 2b**). At d 28 post implantation, histology of tumor xenografts was assessed (**Fig. 2c-h**). CNTNAP4 immunostaining was first performed in xenograft tumors and confirmed a 91.0% reduction in staining among *CNTNAP4* KO implants (**Fig. 2c, d**). Immunofluorescent staining for Ki67 demonstrated a reduction in the proliferative index of *CNTNAP4* KO implants when compared to VC implants (**Fig. 2e, f**, 55.2% reduction in Ki67 labeling). Tumor associated angiogenesis also showed a significant reduction among *CNTNAP4* KO implants compared to control (**Fig. 2g, h**, 81.5% reduction in CD31 labeling), in line with prior reports of the angiogenic effects of CNTNAP4 signaling ^11^. In addition, vascular histomorphometric analysis confirmed that *CNTNAP4* KO had significantly reduced vessel percentage (51.1%), reduced branching index (71.0%), reduced total vessel length (59.0%) (**Supplementary Figure 5**). We further investigated the invasion and metastatic potential in relation to *CNTNAP4* KO (**Fig. 2i, j**). Pulmonary metastases were assessed by immunostaining of human nuclear marker (HuNu) on serial cross-sections of lung tissue, at 28 d post tumor implantation (**Fig. 2i**). We observed pulmonary metastasis in 11/12 (91.7%) of mice with control tumors, in comparison to 5/12 (41.7%) of mice with *CNTNAP4* KO tumors (Chi-square, *p*=0.0094). Overall pulmonary burden of disease was quantified, by assessing the overall number of HuNu positive cells per cross-section of lung tissue. A conspicuous 76.9% decrease in pulmonary HuNu^+^ cell number was observed among mice *CNTNAP4* KO implanted when compared to control among all 12 samples (**Fig. 2j**). Together, our data demonstrate that *CNTNAP4* gene deletion slows OS tumor growth and reduces metastatic spread in an orthotopic xenograft model.

**Figure 2.**
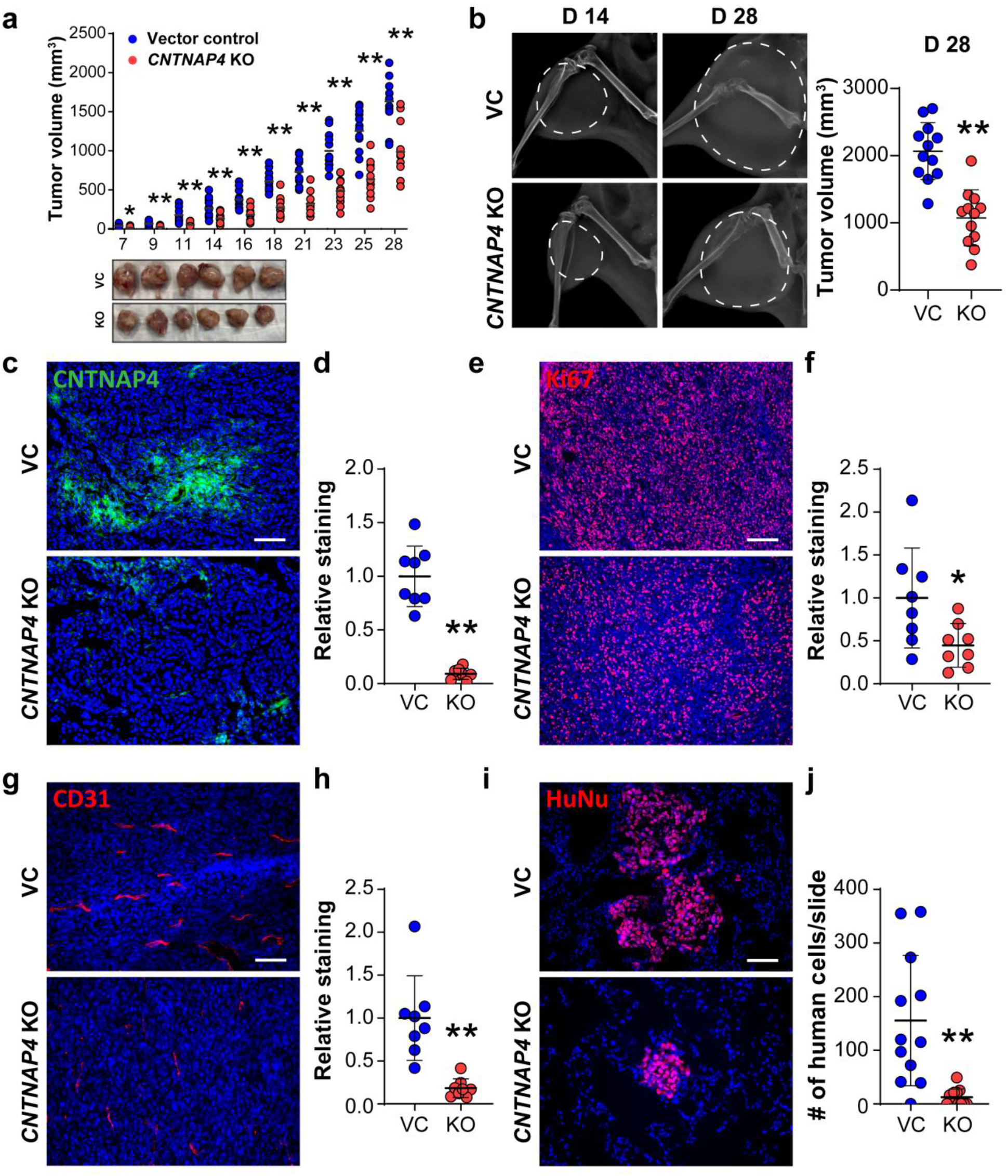
*CNTNAP4* knockout mitigates OS disease progression in 143B xenograft model. Orthotopic implantation of *CNTNAP4* KO or vector control clonal 143B cells within the proximal tibia of female NOD-Scid mice (n=12 mice per group, 1 × 10^6^ cells per implant). (**a**) Tumor volume, calculated by caliper measurements twice weekly until 28 d post-injection (top) and gross pathology of tumors (bottom). (**b**) Representative XR imaging at 14 and 28 d post-injection (left) and tumor volume (d28, right). White dashed line indicating tumors. (**c, d**) *CNTNAP4* KO verified by (**c**) CNTNAP4 immunostaining and (**d**) quantification. (**e, f**) *In vivo* tumor proliferation assessed by (**e**) Ki67 immunostaining and (**f**) quantification. (**g, h**) Tumor associated angiogenesis assessed by (**g**) CD31 immunostaining and (**h**) quantification. (**i, j**) Lung metastasis assessed by (**i**) Human Nuclei (HuNu) immunostaining on cross-sections of pulmonary fields and (**j**) quantification of metastatic cells. Data shown as mean ± 1 SD, with dots representing individual data points. *P*<0.05; \*\**P*<0.01. Scale bars: 100 µm.

### Transcriptomic and proteomic profile after *CNTNAP4* gene deletion

To characterize the transcriptomic landscape following *CNTNAP4* gene deletion in 143B cells, RNA-seq analysis was performed (**Fig. 3**). PCA and differential expression revealed that the *CNTNAP4* KO groups exhibited significantly different expression patterns compared to VC (**Supplementary Figure 6a, b)**. 8,993 DEGs were identified (3,472 upregulated genes and 5,521 downregulated genes) relative to VC (FDR<0.05) (**Fig. 3a**). Top 200 DEGs are provided in **Supplementary File**. Subsequently, IPA and GO term analyses were performed. In *CNTNAP4* KO cells, IPA showed downregulation of synaptic function, a known role of Cntnap4 in neurons (**Fig. 3b**). Consistent with our *in vitro* findings, functional GO enrichment analysis showed downregulation of DEGs in biological processes related to cellular division, cell migration, and cell-cell adhesion, (**Fig. 3c**) as well as other signaling pathways generally related to cancer (**Supplementary Figure 6c, d)**. *CNTNAP4* KO efficiency was further verified by looking at certain key target genes (**Fig. 3d**; *MAST3, CASK, TIAM1, NRX2, NRX3)* that are known to physically interact with *CNTNAP4* (**Supplementary Figure 7**). In addition, we observed the downregulation of key human OS related genes with specific functions in cell cycle/apoptosis (*MDM2, TP53AIP1, RB1-DT*) and DNA repair/damage (*BRCA1, BRCA2, ATM, WRN*) (**Fig. 3e**). Altogether, our transcriptomic analysis corroborated key functional changes with *CNTNAP4* gene deletion and identified enriched molecular targets in the progression of OS.

**Figure 3.**
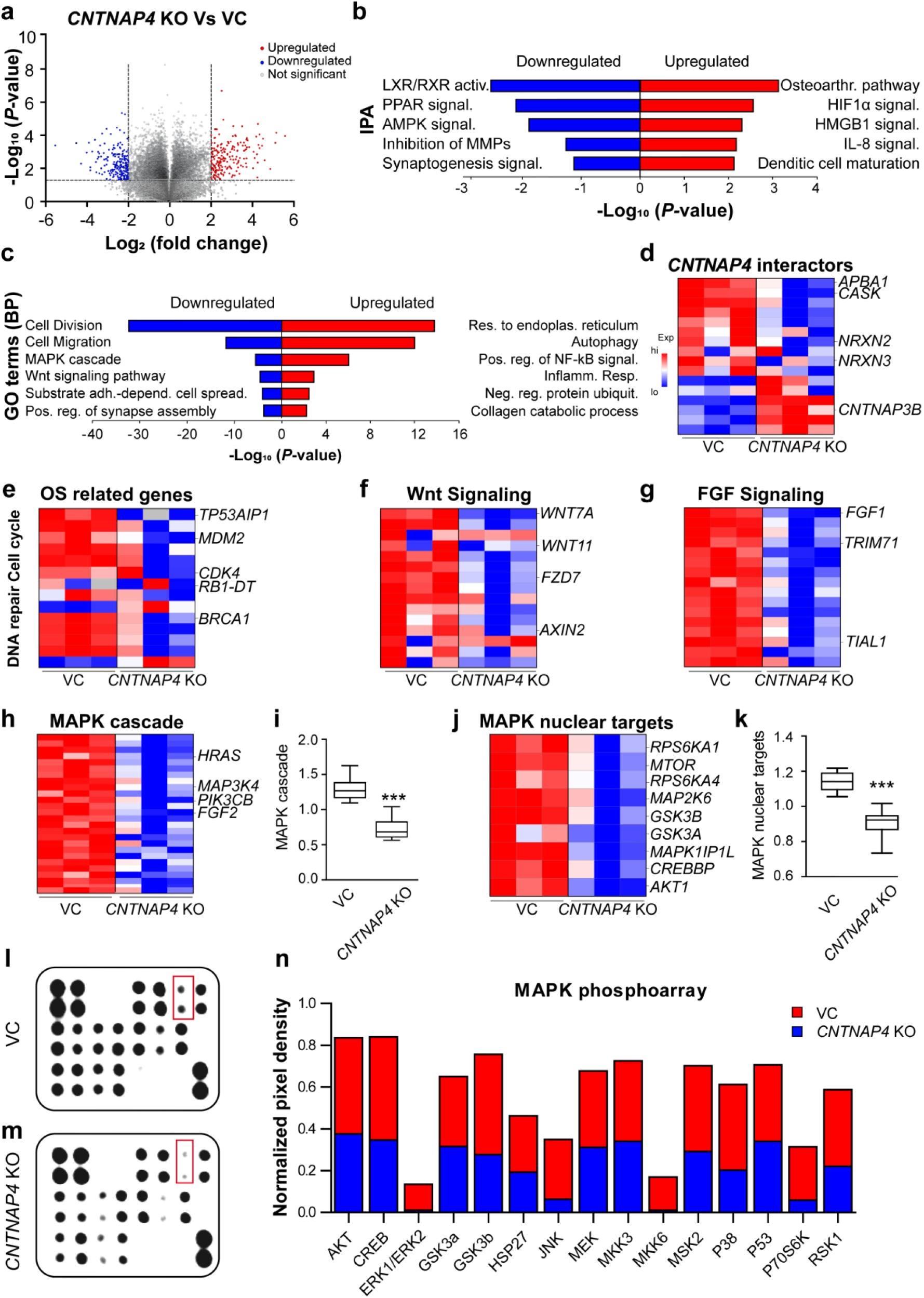
Bulk RNA sequencing between *CNTNAP4* vector control (VC) and knockout (KO) 143B osteosarcoma (OS) cells. **(a)** Volcano plot summarizing RNA-Seq specific differentially expressed genes (DEGs) (FDR <0.05). The red dots on the top right quadrant are significantly up-regulated DEGs and the blue dots within the top left quadrant show highly down-regulated DEGs in the *CNTNAP4* KO compared to VC; grey dots denote unchanged genes. Values are presented as the log2 of fold change. n=3 biological replicates of VC and *CNTNAP4* KO cells. **(b)** Ingenuity Pathway Analysis (IPA) of the RNAseq showing the top canonical pathways enriched in the *CNTNAP4* KO compared to VC. **(c)** Gene Ontology (GO) analysis of RNA-seq shows the classification of the entire transcriptome in biological process (BP). **(d)** Heatmap of *CNTNAP4* interactors that are differentially expressed in KO group. Note that *CNTNAP4* KO decreased expression level of nuclear targets *APBA1*, *CASK* and neurexin family (*NRX2*, *NRX3*). **(e)** Heatmap showing downregulation of OS related genes involved in DNA repair and cell cycle control. Note that *CNTNAP4* KO decreased the expression level of nuclear targets *TP53AIP1*, *MDM2*, and *BRCA1*. **(f)** Heatmap of representative Wnt signaling pathway related genes. Note that *CNTNAP4* KO decreased expression level of certain key genes (*WNT7A*, WNT11, *FZD7, AXIN2*) with implications in cancer progression. **(g)** Heatmap of representative fibroblast receptor growth factor (FGF) signaling. Note that *CNTNAP4* KO decreased expression level of certain key genes (*FGF1*, *TRIM71*, *TIAL1*) with implications in cancer progression. **(h)** Heatmap of representative MAPK signaling pathway related genes showing decreased expression level in *CNTNAP4* KO. **(i)** Box plot showing the average of fold change values from all target genes in the heatmap. **(j)** Heatmap of specific MAPK nuclear targets related genes showing decreased expression level in *CNTNAP4* KO. **(k)** Box plot showing the average of fold change values from all target genes in the heatmap. MAPK phospho-array in *CNTNAP4* VC and KO cells. The human phospho-MAPK array was used to detect eighteen phosphorylated kinases in **(l)** *CNTNAP4* VC and **(m)** *CNTNAP4* KO. Red box indicates the changes in the ERK1/2 spots. **(n)** Stacked column charts depict the mean pixel density of protein levels in lysates prepared from VC (red) and KO (blue). Boxplot shows center line as the median, box limits as upper and lower quartiles of the modulus score. \*\*\**P*<0.001. A two-tailed student’s t test was used for comparisons between *CNTNAP4* KO vs VC.

To better understand the biological function of these differentially expressed mRNAs, GO term analysis was performed on all target genes of deregulated mRNAs. All target genes were enriched into seventeen signaling pathways (**Supplementary Table 1**), including Wnt signaling pathway (53 genes; **Fig. 3f**), Fibroblast growth receptor (FGFR) signaling pathway (27 genes; **Fig. 3g**), MAPK cascade signaling pathway (90 genes). Heatmap and module scoring depicts significantly downregulated genes in the KO group as compared to VC and highlights the certain key genes involved in MAPK signaling, such as *HRAS, MAP3K4, PIK3CB,* and *FGF2* **(Fig. 3h, I, Supplementary Table 2)**. Based on this, we further narrowed our focus to assess the expression of MAPK downstream targets (*MTOR, GSK3A, MAPK1, AKT1, CREBBP, RPS6KA1, RPS6KA4*) which were found to be downregulated in the KO group as depicted in the heatmap (**Fig. 3j**) and box plot (**Fig. 3k**; *p*=0.001). To further validate the downregulation of the MAPK signaling pathway, the levels of phosphorylated MAPK proteins in 143B cells with or without *CNTNAP4* gene deletion were assayed using a Phospho-MAPK Antibody Array **(Fig, 3l, m)**. Overall, there was downregulation of all the MAPK proteins following *CNTNAP4* deletion. The maximum fold reduction was observed in ERK1/2 (12.5-fold), JNK (4.45-fold) and RPS6KB1 (4.32-fold) (**Fig. 3n**). Thus, a transcriptomic-phosphorylation array analysis confirmed that *CNTNAP4* deletion significantly reduces MAPK signaling activity, including prominent reductions in ERK1/2 signaling activity.

### ERK1/2 agonist reverses effects of *CNTNAP4* deletion

Having observed a prominent reduction in ERK1/2 signaling in *CNTNAP4* KO cells, we next examined whether an ERK agonist could restore MAPK signaling and cellular behavior among *CNTNAP4* KO 143B cells. Here, either VC or *CNTNAP4* KO 143B cells were treated with the ERK1/2 agonist isoproterenol (ISO)^12^, and proliferation, attachment, migration, and invasion were again assayed (**Fig. 4a-d**). When treated with ISO, *CNTNAP4* KO cells showed a complete or near complete restoration of cellular proliferation to levels comparable to VC cells (**Fig. 4a**). Similar observations were made with ISO-treated *CNTNAP4* KO cells in rates of cell attachment, migration, and invasion (**Fig. 4b-d**). Furthermore, phospho-array confirmed a restoration of pERK1/2 (7.34-fold increase) and other MAPK associated proteins among ISO-treated *CNTNAP4* KO cells such as JNK and p38 **(Fig. 4e, f)**. Western blot further confirmed the restoration of pERK1/2 and pJNK in ISO-treated *CNTNAP4* KO cells (**Fig. 4g**, **Supplementary Figure 8**). The activation of ERK1/2 signaling was next assessed by immunofluorescent staining for ERK1/2 and pERK1/2 across 143B xenografts. Among *CNTNAP4* KO tissue sections, decreased pERK1/2 and retained ERK1/2 immunoreactivity was observed in comparison with control tumors (**Fig. 4g**).

**Figure 4.**
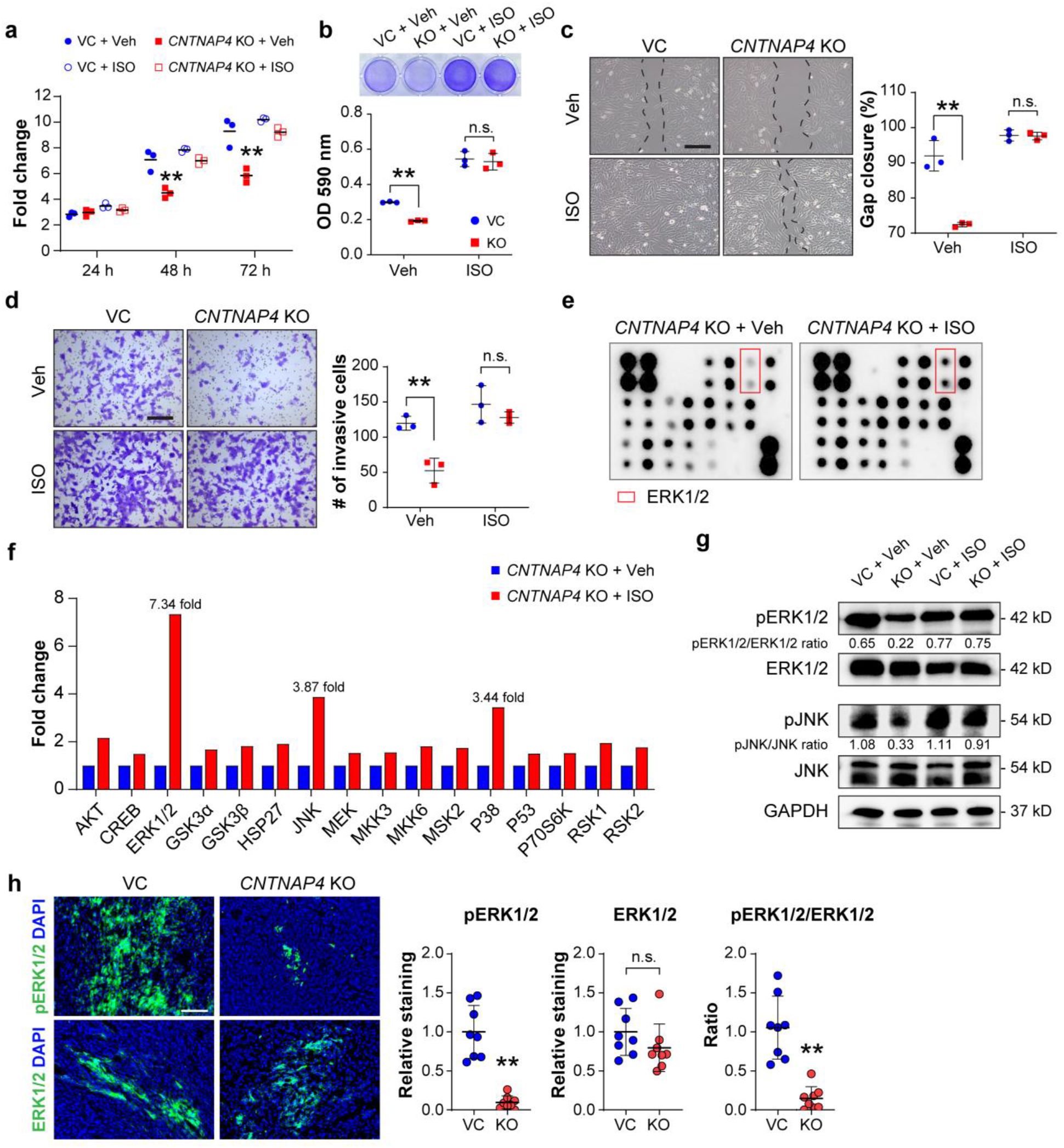
ERK agonist restores cellular phenotypes in *CNTNAP4* KO cells. Cells were pre-treated by vehicle control (Veh) or 1 μM isoproterenol (ISO) for 5 min. (**a**) MTS proliferation assay among cells at 24, 48, and 72 h. (**b**) Attachment assay at 3 h, assessed by crystal violet staining (top) and quantification (bottom). (**b** Attachment assay at 3 h, assessed by crystal violet staining (top) and quantification (bottom). (**c**) Migration assay at 24 h assessed by scratch wound healing assay (left) and quantification (right). (**d**) Transwell invasion assay at 24 h, including representative images (left) and quantification (right). Data shown as mean ± 1 SD, with dots representing individual data points. \*\**P*<0.01; n.s.= not significant. Scale bars: 100 µm. (**e**) Protein array demonstrating MAPK phosphorylation in *CNTNAP4* KO clones with or without ISO treatment. (**f**) Phosphorylation level of MAPK proteins measured as fold-change of the paired duplicate spots and normalized to positive control spots. (**g**) Western blot of pERK1/2, ERK1/2, pJNK and JNK. Ratio of pERK1/2/ERK1/2 and pJNK/JNK is listed below. (**h**) pERK1/2 and ERK1/2 immunostaining and quantification in 143B xenografts.

### *CNTNAP4* KO and *NELL1* KO sarcoma cells display similarities and differences in transcriptomic profiling

Thus far, significant similarities had been observed between *CNTNAP4* KO sarcoma cells and those effects we recently reported in *NELL1* KO cells ^6^. To further compare 143B sarcoma cells with *NELL1* or *CNTNAP4* gene deletion, direct transcriptomic pathway analysis was performed (**Fig. 5**). Analysis of GO terms showed some clear similarities and differences between datasets including enrichment for terms such as axon guidance, cell adhesion, cell proliferation, and cell migration with either *CNTNAP4* or *NELL1* gene deletion (**Fig. 5a**). Notably, GO term enrichment predicted a larger role for cell division with *CNTNAP4* KO as opposed to *NELL1* KO when compared to their respective controls. The transcriptional similarities and differences between *CNTNAP4* and *NELL1* gene deletion were next assessed by KEGG pathway analysis (**Fig. 5b**). Using significantly downregulated and upregulated KEGG pathways (p<0.05), Venn diagrams showed only 22 shared down-regulated pathways and 18 shared up-regulated signaling pathways between *CNTNAP4* and *NELL1* KO datasets (**Fig. 5c, Supplementary File**). Further analysis of each of the downstream pathway presented the gene expressions are distinctly different between *CNTNAP4* KO and *NELL1* KO. *CNTNAP4* target gene expression was next compared across datasets, shown as either heatmap or gene module scores in comparison to each respective control (**Fig. 5d**). Overall, a significant downregulation of *CNTNAP4* target genes was observed in *CNTNAP4* KO cells, but not in *NELL1* KO cells (**Fig. 5d**). Similarly, MAPK and FAK signaling pathways were significantly downregulated in *CNTNAP4* KO cell clones, but to a lesser extent in *NELL1* KO tumor cells (**Fig. 5e, f**). Interestingly, ECM associated genes (previously reported to be significantly reduced in expression with *NELL1* KO) ^6^, were significantly upregulated among *CNTNAP4* KO tumor cells (**Fig. 5g**). These data suggested that *CNTNAP4* KO demonstrate partial overlap in transcriptomic profiling with *NELL1* gene deletion tumor cells, but that crucial differences in MAPK/FAK signaling and ECM expression profiles exist between the two KO transcriptomic phenotypes.

**Figure 5.**
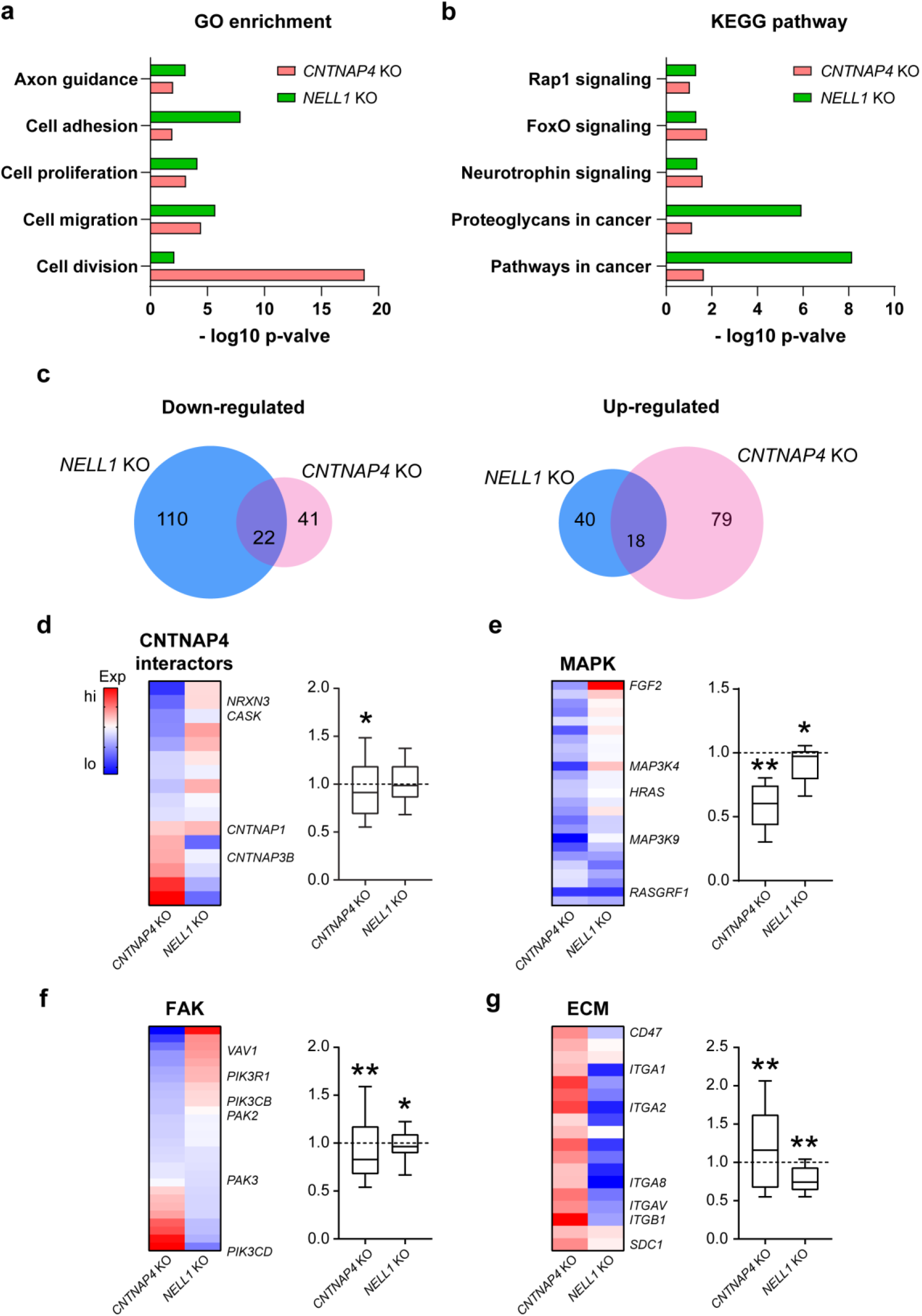
Comparison of bulk RNA sequencing among *CNTNAP4* KO and *NELL1* KO OS cells. (**a-f**) Total RNA sequencing comparison between *CNTNAP4* KO and *NELL1* KO clonal 143B cells. (**a**) Gene Ontology (GO) enriched in both *CNTNAP4* KO and *NELL1* KO cells. (**b**) KEGG pathways enriched in both *CNTNAP4* KO and *NELL1 KO* cells. (**c**) Venn diagrams show the similarities and differences in the significantly up- and down-regulated pathways between *NELL1* KO (blue) and *CNTNAP4* KO (pink) datasets (*P*<0.05). (**d-g**) Heat map demonstrating the contrast in the expression levels of genes involved in downstream signaling and corresponding module score among *CNTNAP4* KO and *NELL1* KO cells, including (**d**) *CNTNAP4* interactors, (**e**) MAPK signaling, (**f**) FAK signaling, and (**g**) ECM-receptor interaction. Gene module scores are shown as a boxplot shows center line as the median, box limits as upper and lower quartiles of the modulus score. The dashed line indicates no change in gene module score in comparison to vector control. \**P*<0.05; \*\**P*<0.01, in comparison to the corresponding vector control.

### Clinical relevance of *CNTNAP4* in human osteosarcoma

First, expression of Cntnap4 was verified in different sets of human osteosarcoma tissues and primary cells. Immunohistochemical staining of primary OS resection tissue sections with viable tumor showed consistent immunoreactivity across all tumor samples (**Fig. 6a, b,** N=8). Likewise, qRT-PCR demonstrated amplification of *CNTNAP4* mRNA across all human OS primary cells examined (**Fig. 6c**, N=4). Next, we reanalyzed a scRNA-Seq dataset which utilized a patient-derived xenograft model of osteosarcoma ^13^. Confirming the author’s original analysis, we identified both *Ki67*^high^ and *Ki67*^low^ sarcoma cells (**Fig. 6d**). Although the expression of *CNTNAP4* in this dataset had lowly expressed transcripts, the levels of *CNTNAP4* interacting genes (*MACF1, MLLT4, FBXO21, CASK*) were found to be highly enriched in the *Ki67*^high^ sarcoma cells (**Fig. 6d**), which was found as well to cluster with high *MDM2* expression. To investigate whether *CNTNAP4* expression correlates with prognosis in OS patients, we performed a meta-analysis on public gene-expression data from human soft tissue sarcoma dataset and obtained a graphical summary of genomic alterations in multiple *CNTNAP4* interacting genes (**Fig. 6e**). The number of patients and frequency of genetic alteration in each tumor sub-type is provided in **Supplementary Figure 9**. Among the queried genes, we found maximum genetic alterations in *CNTNAP3B* (8%), *APBA1* (6%), *CNTNAP2* (6%), *NELL1* (4%), *CASK* (3%), *TIAM1* (3%), and *NRX3* (2.9%). Most of the genes showed amplifications. A total of seventy cases showed alterations in one of the query genes with thirty-two events (median survival – 54.20 months) while the group without genetic alterations had 136 cases with 46 events (median survival 80.50 months). Notably, patients with genetic alterations in *CNTNAP4* associated genes had shorter overall survival than those without alterations (log rank, *p*=0.04; **Fig. 6f**).

**Figure 6.**
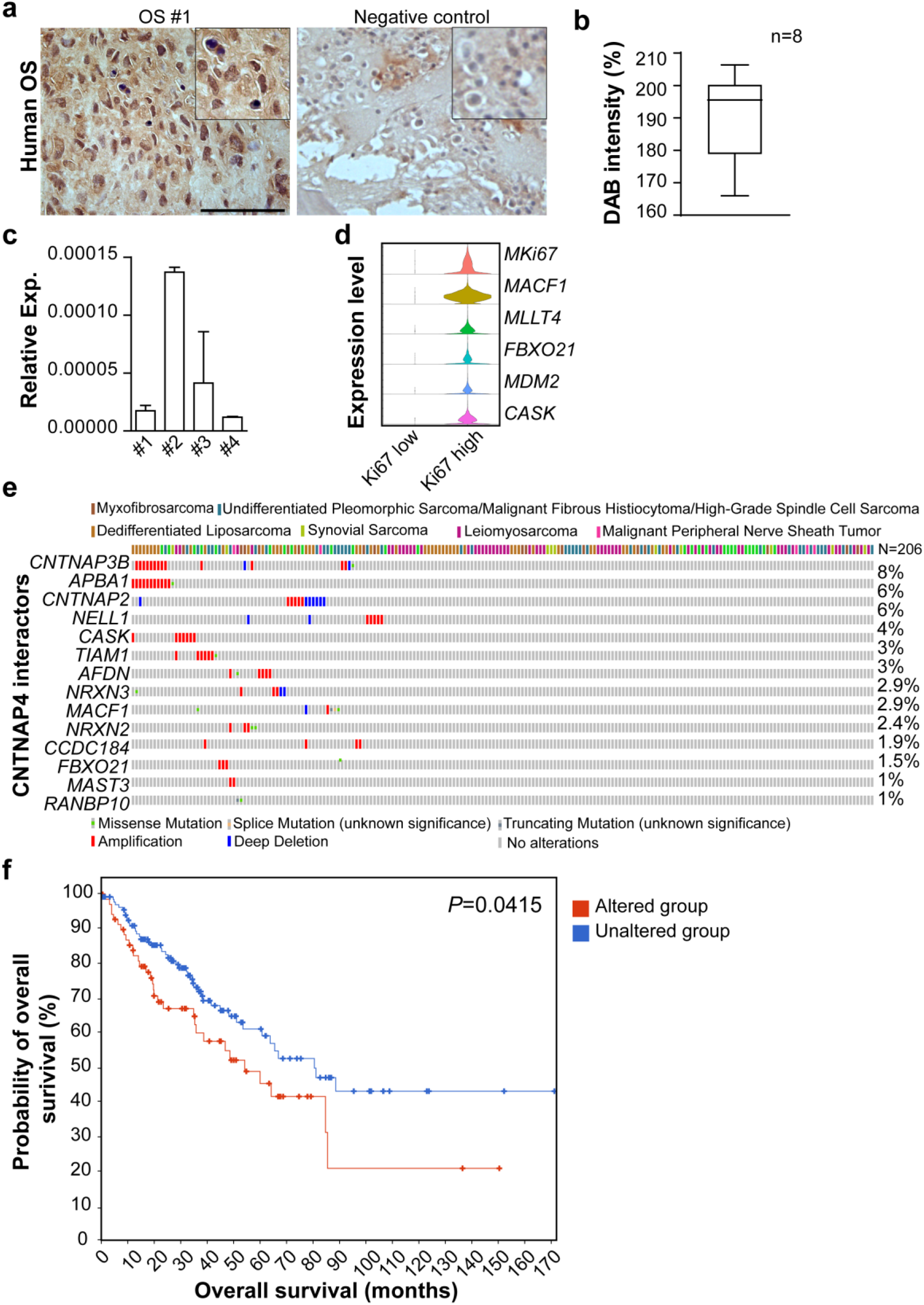
Clinical relevance of Cntnap4 and other interactors in Osteosarcoma (OS). **(a)** Representative immunohistochemical staining of CNTNAP4 within human OS tissue section (left) and negative control (right); high magnification is provided as inset. Scale bar: 100 µm. **(b)** Quantification of DAB intensity in human OS tissue sections (n=8). **(c)** Relative *CNTNAP4* mRNA expression in primary human OS cells. **(d)** Single cell RNA-Seq analysis of patient-derived xenograft OS sample. Violin plots show gene expression of *CNTNAP4* interacting genes (*MACF1, MLLT4, FBX021, CASK*) in the two clusters, *KI67*^low^ (101 cells) and *KI67* ^high^ (3,645 cells). **(e)** Genomic alterations in samples from an Adult Soft Tissue Sarcoma (TCGA, Cell 2017) dataset from cBioPortal. Waterfall plot depicts the genetic alterations in *CNTNAP4* interacting genes across multiple sarcoma subtypes. Red represents amplification and blue represents deep deletion of genes in 34% of the sarcoma patients (n=206). **(f)** Kaplan-Meier curve between groups with and without genetic alterations in at least one of the 15 genes including *CNTNAP4* (*p*=0.0415). The red and blue lines represent cases with and without genetic alterations, respectively. Data shown as mean +1 SD. Boxplot shows center line as the median, box limits as upper and lower quartiles of the DAB intensity.

## Discussion

In summary, we report that *CNTNAP4* gene deletion mitigates aggressive sarcoma behavior in an orthotopic model of human OS, characterized by delayed tumor onset, decreased tumor progression, reduced tumor-associated angiogenesis and diminished ERK/MAPK signaling. This was corroborated by several cellular changes including reduced cell migration, invasion, and altered transcriptional targets of ERK signaling pathway. Cntnap4 exerts its effects partly via interaction with NELL-1, an osteosarcoma associated protein, and subsequent positive regulation of MAPK intracellular components. Finally, we confirmed that the diminished ERK1/2 activity due to loss of *CNTNAP4* can be restored by an ERK agonist. We conclude that NELL1/CNTNAP4 activates ERK/MAPK signaling, and this is a feature of an aggressive phenotype in osteosarcoma (**Fig. 7**). Further, perturbing NELL-1/CNTNAP4 signaling could serve as a potential therapy to improve OS disease progression.

**Figure 7.**
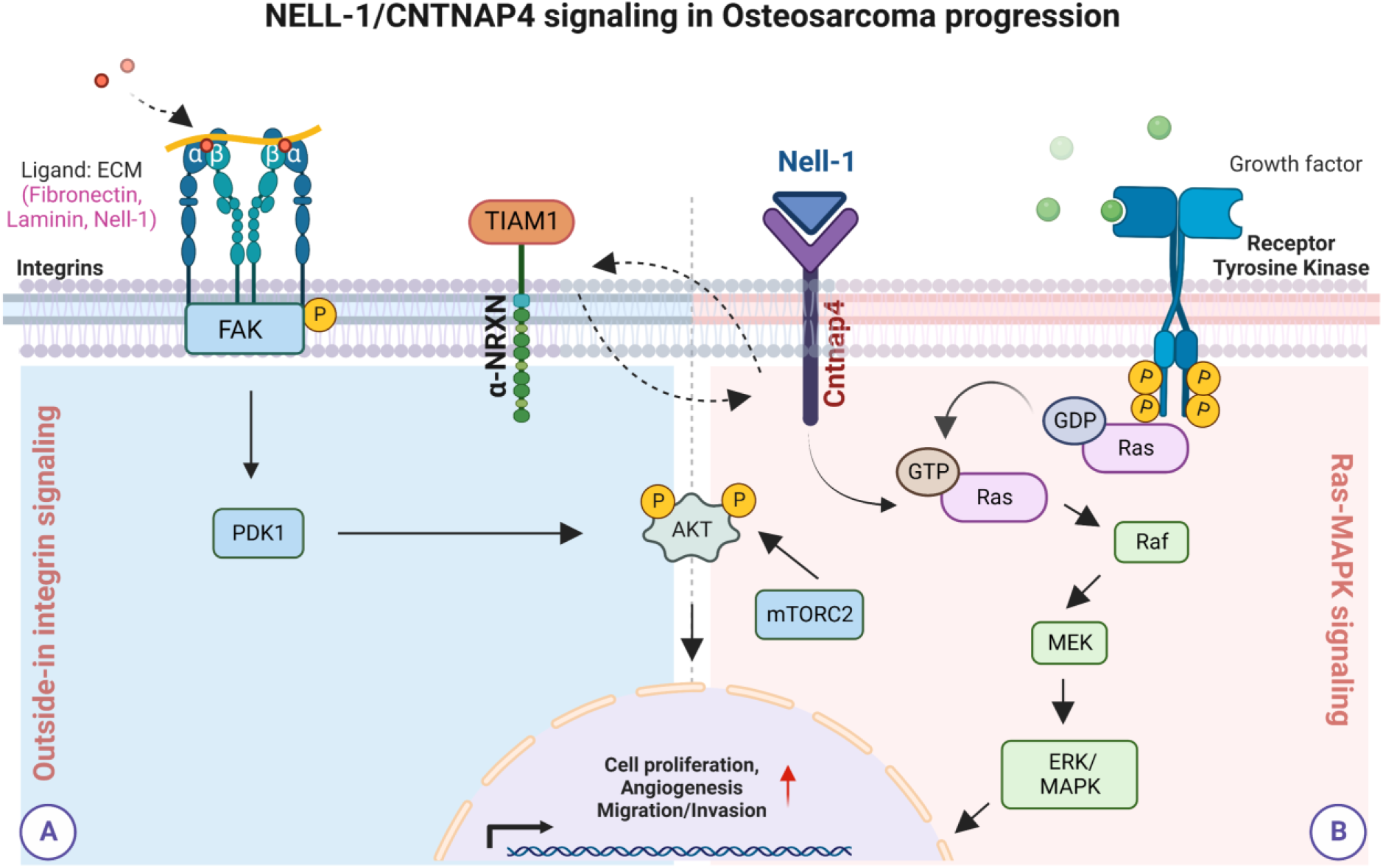
Model proposing mechanism driving sarcoma progression through Nell-1/Cntnap4 signaling. As a secreted molecule, NELL-1 initiates cellular signaling through binding to its receptor, Cntnap4, on the cell surface. The Ras-MAPK/ERK and Outside-in integrin signaling pathways play critical roles initiating downstream targets during sarcoma disease progression. Cartoon created with BioRender.com.

We previously reported that NELL-1, a protein known to regulate osteogenesis ^14^, has a positive impact in OS disease progression ^5, 6^. NELL1 modulates the sarcoma matrisome and cell-ECM interactions to promote sarcomagenesis and disease progression. We have now extended this work to elucidate the role of CNTNAP4, a high-affinity receptor for NELL-1 in mediating tumor progression. Significantly, several genetic alterations that collectively interact with *CNTNAP4* have poor prognosis in patients with soft tissue sarcoma, although no data for osteosarcoma is yet known.

Prior studies have demonstrated that Cntnap4 has a distinct pattern of cellular localization from that of other members in the family (Cntnap, Cntnap2, Cntnap3, Cntnap5) ^15^, indicating that they may have diverse functions inside and outside of central nervous system ^8^. By structural homology, Cntnap4 most closely resembles Cntnap3 followed by Cntnap2, Cntnap and neurexins ^15, 16^. Owing to this structural and functional similarity it is worthwhile to speculate the contribution of other Cntnap subunits to balance the deficit of Cntnap4 function in our study. Despite their high structural similarity, the extracellular domain of the Cntnap4 facilitate interactions with multiple ligands that are members of the Contactin subgroup (TAG-1, BIG-1, BIG-2, NB2, NB3, and FAR-2) to form cell-recognition complexes ^15^. On the other hand, the cytoplasmic region of Cntnap4 (C4ICD) comprises a carboxyl-terminal binding site exclusive for binding different PDZ domain-containing proteins involved in regulating cellular and biological processes ^17^. Some of the binding partners of Cntnap4 that are predicted to localize at plasma membrane, cytoplasm, and nuclear include CASK, TIAM1, APBA1 (MINT1), LNX2, MAST3, NRXN, MACF1. Interestingly, overexpression of these regulatory proteins such as CASK, TIAM1, APBA1, and NRXN, have also been linked with sarcoma progression ^18, 19^.

While studies reveal high-affinity interaction between N-terminus of NELL-1 and Cntnap4 ^20^, several other cell-surface membrane receptors that physically interacts with NELL-1 via N-terminal thrombospondin-1 (TSP-N) and epidermal growth factor (EGF) domains have been identified to facilitate its downstream signaling cascade during neurogenesis, osteogenesis, chondrogenesis, vasculogenesis, and sarcomagenesis ^20–23^. Some of the other physical protein interactions of NELL-1 include integrin β1 (ITGB1), protein kinase C β1, apoptosis-related protein 3 (APR3) and Roundabout 2 (Robo2) ^10, 24–26^. This suggests that NELL-1 exerts at least some effects independent of Cntnap4 and that NELL-1 could have more diverse effects outside of Cntnap4-MAPK-FAK signaling in sarcoma biology. Nevertheless, we observed under certain doses of NELL1 protein and in the assayed examined that *CNTNAP4* KO cells remained unresponsive to NELL1 treatment. Therefore, we cannot determine the extent to which other NELL-1 binding partners play a role in NELL-1 mediated changes in sarcoma cell behavior ^10^.

There are some limitations to the present study. First, the observed human genetic data in soft tissue sarcomas represented several distinct sarcoma types and different genetic alterations, including both loss-of-function and gain-of-function mutations. Therefore, future studies investigating the role of NELL-1/CNTNAP4 function in sarcoma progression will likely require prospective data collection on single tumor types, including bone-associated sarcomas. Second, it is clear, as alluded to, that both NELL-1 and CNTNAP4 have several binding partners that may play partially redundant roles in sarcoma disease progression. Use of additional genetic deletions, such as combination NELL-1/CNTNAP4 knockout alleles, will assist in further decoding the effects of these protein targets in OS biology. Third, this study has mechanistically focused on the changes primarily associated with the MAPK/ERK pathway following *CNTNAP4* deletion. We recognize many other transcription factors and indeed changes mediated by Isoproterenol affects multiple downstream events. Further studies may be necessary to evaluate the principal changes associated upstream or downstream of ERK1/2 such as PKA/CREB. Taken together, our results demonstrate that CNTNAP4 signaling plays an important regulatory role in OS disease progression. These findings suggest that targeted treatment of the Nell-1/Cntnap4 signaling axis represents a potential strategy to mitigate sarcoma disease progression.

## Methods

### Mice

All animal experiments were conducted according to approved protocols (MO21M112) of the Animal Care and Use Committee (ACUC) at Johns Hopkins University (JHU). NOD Scid mice (Stock No: 001303) were procured from The Jackson Laboratory (Bar Harbor, ME, USA).

### Cell isolation and culture

Human OS cell lines were procured from American Type Culture Collection (Manassas, VA), including 143B (ATCC®-CRL-8303™), Saos-2 (ATCC^®^ HTB-85^™^), HOS (ATCC CRL-1543), KHOS/NP (ATCC CRL-1544), KHOS-312H (ATCC CRL-1546), and G-292 (ATCC CRL-1423). Primary human OS cells were harvested from human OS resection samples (n=4) under IRB approval at JHU. The diagnosis of high-grade conventional OS was verified by two independent bone pathologists (E.F.M. and A.W.J.). The human OS tissue samples collected under sterile conditions were washed in PBS and dissected into small bits (<1 mm^3^). Primary OS tumor cells were isolated as previously described ^27^ and were culture-expanded for 3 to 5 passages prior to use. Patient-derived primary OS cells verified by karyotyping, immunophenotyping (CD31^-^CD45^-^CD44^+^CD73^+^CD90^+^CD105^+^), and multi-lineage differentiation were used ^6, 28^. The primary OS cells were cultured in Dulbecco’s Modified Eagle Medium (DMEM, Gibco, Grand Island, NY) supplemented with 15% fetal bovine serum (FBS, Gibco), 100 U/ml penicillin, and 100 µg/ml streptomycin (Gibco) in a humidified incubator with 5% CO2 at 37°C.

### CNTNAP4 gene deletion and isolation of cell clones

CRISPR/Cas9 gene deletion of *CNTNAP4* in human OS cell lines and primary cells was achieved using a plasmid with a guide RNA (gRNA) sequence combined with U6gRNA-Cas9-2A-RFP vector (Sigma-Aldrich). The specific gRNA sequence of KO and negative control plasmids used in this study is shown in **Supplementary Table 3**. Transient plasmid DNA transfection was performed using TransIT®-LT1 Transfection Reagent (Mirus, Madison, WI) ^6^. Briefly, cells plated at density of 3×10^5^ cells/ml in 6-well cell culture plates were transfected with TransIT-LT1 Reagent-plasmid DNA complex and incubated for 72 h. Next, *CNTNAP4* KO single cell colonies were established by FACS. Viable RFP-positive cells from *CNTNAP4* KO-transfected cells and control-transfected cells were sorted into wells of a 96-well microtiter plate by using a Dako Cytomation MoFlo cell-sorter (Beckman, Indianapolis, IN). For single cell sorting, a 70-mm nozzle with the sheath pressure set at 70 PSI was used. After confirmation of *CNTNAP4* gene deletion by qRT-PCR, n=2 colonies were expanded for use.

### Small interfering RNA (siRNA) and Transfection

Knockdown of *CNTNAP4* in sarcoma cells was performed using Silencer Select chemically synthesized siRNA (Thermo Fisher Scientific, Cat# 4390846; s40010) ^6^. Briefly, cells were seeded in 6-well culture plates at a density of 3× 10^4^ cells per well. At 70% confluence, basal medium was removed and replenished with antibiotic-free basal medium. Transfection was performed using TransIT®-LT1 Dynamic Delivery System (Mirus, Madison, WI) and 100 nm CNTNAP4 siRNA or scramble siRNA diluted in minimal essential medium (Opti-MEM) and incubated for 3 h. *CNTNAP4* knockdown efficiency was confirmed by qRT-PCR at 48h post-transfection.

### T7 Endonuclease I Assay

*CNTNAP4* KO and control-transfected cells were collected for the detection of the mutation using the T7 Endonuclease I Assay Kit (Genecopoeia, Rockville, MD). DNA was extracted from the samples using Quick-DNA™ Miniprep Kit (Zymo Research, Irvine, CA). See **Supplementary Table 3** for primer information. PCR reactions were performed according to the manufacturer’s protocol ^6^. Briefly, the PCR product was digested using T7E1 enzyme at 37°C for 60 minutes, followed by separation using 1.5% agarose gel electrophoresis.

### In vitro functional assays

The following experiments were performed as described previously ^6, 29^. First, cell proliferation was measured using the CellTiter 96® Aqueous Non-Radioactive MTS Cell Proliferation Assay (Promega, Madison, WI). A total of 2 × 10^3^ cells were plated in 96-well plates and incubated for 24, 48, and 72 h. The optical density at 490 nm was measured using a microplate spectrophotometer (BioTek, Winooski, VT). Second, cell attachment was assayed by crystal violet staining. A total of 2 × 10^5^ cells were plated in 24-well plates and incubated for 1, 3 and 5 h. Briefly, the cells were fixed with 100% methanol and then stained with 0.5% crystal violet solution for 10 min at RT. The crystal violet stain was eluted using 10% acetic acid and the absorbance was measured at 570 nm. In select experiments, culture plates were pre-coated with 5 µg/cm^2^ of recombinant NELL1 (provided by X.Z.). Third, migration assays were conducted using the Culture-Insert 2 Well (Ibidi, Martinsried, Germany). A total of 4 × 10^4^ cells per well were incubated overnight. The following day, insert wells were gently removed, and the plate was replenished with growth medium. After incubation for 12 and 24 h, photographs were taken using an inverted microscope (Olympus, Tokyo, Japan). The diameter of the cell-free gap was determined using ImageJ software (Version 1.49 v, NIH, Bethesda, MD). Fourth, invasion assays were conducted using Corning® BioCoat™ Matrigel® Invasion Chambers (Corning, Bedford, MA). Briefly, 2.5 × 10^5^ cells suspended in serum-free medium were added to the upper chamber, and 500 μl growth medium was added to the lower chamber. After 24 h incubation, cells were fixed and stained with 0.5% crystal violet solution. Photographs of five random areas were taken under an inverted microscope (Olympus) and the number of invaded cells was counted. Finally, in select experiments cells were stimulated with vehicle (Veh) or Isoproterenol (ISO), a β-adrenergic agonist, at 1 µM concentration for 5 min at 37°C. Cell proliferation, attachment, migration, invasion was assessed. Experiments were done in triplicate.

### qRT-PCR

Total RNA was extracted from cultured cells of equal passage number and density using TRIzol^TM^ Reagent (Invitrogen, Carlsbad, CA) according to the manufacturer’s instructions. A total of one μg of total RNA was used for reverse transcription with iScript cDNA synthesis kit (Bio-Rad). Real-time PCR was performed using SYBR Green PCR Master Mix (Thermo Scientific, Waltham, MA). Relative gene expression was calculated using a 2^-ΔΔCt^ method by normalization with GAPDH. Primer sequences are listed in **Supplementary Table 3.**

### Bulk RNA-Seq and data analysis

Gene expression was detected by total RNA sequencing using the Illumina NextSeq 500 platform (Illumina, San Diego, CA). Briefly, total RNA was extracted from clonal 143B cells with or without *CNTNAP4* KO by Trizol (Life Technologies Corporation, Gaithersburg, MD) and three independent RNA samples were prepared. The total RNA samples were sent to the JHMI Deep Sequencing and Microarray core for sequencing. Data analysis was performed using software packages including CLC Genomics Server and Workbench (RRID: SCR_017396 and RRID: SCR_011853), Partek Genomics Suite (RRID: SCR_011860), Spotfire DecisionSite with Functional Genomics (RRID: SCR_008858), and QIAGEN Ingenuity Pathway Analysis (IPA, RRID: SCR_008653). Kyoto Encyclopedia of Genes and Genomes (KEGG) and Gene Ontology (GO) enrichment analysis of differential expression genes (DEGs) were performed in Database for Annotation, Visualization, and Integrated Discovery (DAVID) bioinformatics software.

### cBioPortal for Cancer Genomics repository

The cBioPortal for Cancer Genomics repository was utilized to reanalyze multi-omics data from the Cancer Genome Atlas [Adult Soft Tissue Sarcomas (TCGA, Cell 2017)] ^30, 31^. Genomic and transcriptomic data from 206 patients representing three different sarcoma types (Soft tissue Sarcoma, Uterine Sarcoma and Nerve Sheath Tumor) with seven different cancer subtypes [Leiomyosarcoma (25.7%), Dedifferentiated Liposarcoma (24.3%), Undifferentiated Pleomorphic Sarcoma (21.4%), Uterine Leiomyosarcoma (13.1%), Myxofibrosarcoma (8.3%), Synovial Sarcoma (4.9%), Malignant Peripheral Nerve Sheath Tumor (2.4%)] were analyzed. A list of CNTNAP4 interacting genes (*CNTNAP3B, APBA1, CNTNAP2, NELL1, CASK, TIAM1, AFDN, NRXN3, MACF1, NRXN2, CCDC184, FBXO21, MAST3, RANBP10*) was analyzed. The OncoPrint provided the genetic alteration frequency on the DNA level for the different cancer types for the queried gene list. Genetic alterations were further divided into missense mutations, amplification, deep deletion, and multiple alterations. From the same platform, we obtained the overall patient survival status.

### Single-cell RNA sequencing (scRNA-Seq)

Publicly available scRNA-Seq data was obtained (accession number: GSE179681, Sample GSM5426214) ^13^. The single cell transcriptomic library of 47,977 OS cells was obtained from a patient-derived xenograft (PDX) model, grown as an orthotopic tumor. Seurat R package was used for quality control, filtering, and analysis of gene expression matrices. Cells were filtered to remove doublets, low quality cells, and cells with high mitochondrial genes. Cells with fewer than 100 expressed genes and genes expressed in fewer than five cells were removed. Downstream analysis was performed using Seurat v3.0 package ^13^. Expression matrices were converted to Seurat object through the “CreateSeuratObject” function. Cells that that have unique feature counts over 2,500 or less than 200 were digitally filtered out for further analysis. Next, the dataset was scaled, analyzed for principal components, and visualized using UMAP. Human cells were differentiated by Ki67 expression [cluster 0 (Ki67 low): 101 cells, cluster 1 (Ki67 high): 3,645 cells]. Along with *MDM2*, a negative regulator of the p53 tumor suppressor, expression of genes with predicted physical interactions with CNTNAP4 were analyzed (*MACF1, MLLT4, FBXO21, CASK*).

### MAPK phosphorylation protein array

Protein array analysis was performed using the Human MAPK Phosphorylation Antibody Array (ab211061) according to the manufacturer’s instructions (Abcam, Boston, MA). Visualization was performed with the ChemiDoc Touch Imaging System (Bio-Rad, CA). The average signal (pixel density) of the pair of duplicate spots representing each MAPK downstream protein was normalized on an averaged positive control spot. Corresponding signals were compared to determine the relative difference in MAPK downstream proteins among cells with or without *CNTNAP4* gene deletion. Data shown are from automatic exposure second exposure using a chemiluminescence imaging system. See **Supplementary Table 4** for protein layout.

### Western Blot

Western blot was performed as previously described^6^. Briefly, cells were lysed in RIPA buffer with protease inhibitor cocktail (Cell Signaling Technology), and protein concentrations were determined by BCA assay (Thermo Scientific). Cell lysates were then separated by SDS-PAGE and transferred onto nitrocellulose membrane. Membranes were then blocked with 5% BSA and incubated with primary antibodies at 4 °C overnight. Membranes were incubated with a horseradish peroxidase (HRP)-conjugated secondary antibody and detected with ChemiDoc XRS+ System (Bio-Rad). All western blots within a panel are from the same experiment and all blots were processed in parallel. Antibodies are listed in **Supplementary Table 5**.

### Osteosarcoma implantation and assessments

All animal studies were performed with institutional ACUC approval within Johns Hopkins University. For all 143B OS cell implantation, 8–10-week-old, male and female NOD Scid mice were used. 1 × 10^6^ clonal 143B cells with or without *CNTNAP4* KO in 50 μl PBS were injected subperiosteally within the proximal tibia metaphysis. Tumor size was measured by caliper three times weekly or radiographs every 2 weeks. Tumor formation was monitored and recorded, and tumor volume was calculated ^6^. Tumor volume was calculated using an inbuilt measurement tool in the radiograph software and the tumor size was measured and calculated as per caliper measurement. Primary tumors and lungs were harvested at 28 d post-injection for histologic analysis. Analyses were performed by investigators blinded to the treatment group.

### Histology and Immunohistochemistry

Primary tumor samples and lungs were fixed in 4% paraformaldehyde (PFA) at 4°C for 24 h and decalcified in 14% EDTA (Sigma-Aldrich) for up to 28 d at 4°C. Samples were then cryoprotected in 30% sucrose overnight at 4°C and embedded in optimal cutting temperature compound (OCT, Tissue-Tek 4583, Torrance, CA) to obtain frozen sections. Routine H&E staining was performed on the entire lung tissues (10 µm thick) sectioned in a coronal plane to assess pulmonary burden, and number of metastasis foci were manually counted. For immunohistochemistry, sagittal sections of primary or metastatic tumors were washed in PBS × 3 for 10 min and permeabilized with 0.5% Triton-X for 30 min. Next, 5% normal goat serum (S-1000, Vector Laboratories, Burlingame, CA) was applied for 30 min, then incubated in primary antibodies overnight at 4°C. The following day, slides were washed in PBS, incubated in the secondary antibody for 1 h at RT, and then mounted with DAPI mounting solution (Vectashield H-1500, Vector Laboratories). Digital images of these sections were captured with 10-100 × objectives using upright fluorescent microscopy (Leica DM6, Leica Microsystems Inc., Buffalo Grove, IL). Analyses were performed by investigators blinded to the sample identification. Antibodies are listed in **Supplementary Table 5**.

### Vascular morphometry

For the quantitative vascular morphometric analysis, a semi-automated, validated, open-source software (Angiotool 0.6a, http://angiotool.nci.nih.gov) was utilized to quantify the differences in morphological and spatial parameters of the vascular network between *CNTNAP4* VC (n=4) and KO (n=8) tumor xenografts. The following angiogenesis related parameters were assessed in the CD31 immunoassay samples: vessel percentage, total vessel length, junction density and total number of endpoints.

### CNTNAP4 expression in human osteosarcoma

Human OS resection tumor sections (n=8) were used under IRB approval at JHU with a written informed consent for tissue banking. Paraffin-embedded tissue sections were deparaffinized in xylene and rehydrated in descending grades of alcohol solutions. Heat-induced epitope retrieval was performed using 10 mM sodium citrate buffer, 0.05% Tween 20 (pH 6.0) for 20 min at 85-90°C. Endogenous peroxidase activity was blocked using BLOXALL™ Blocking Solution (Vector laboratories, Burlingame, CA, USA) for 10 min. Sections were washed in 1x PBS and incubated overnight at 4°C in humid chambers with anti-CNTNAP4 primary antibody (Biorbyt, United Kingdom). The following day, sections were washed in 1x PBST (3x) and incubated with a secondary antibody (Invitrogen, Waltham, MA) for 1 h at RT. The sections were further incubated with avidin-biotin complex (Vector laboratories, Burlingame, CA, USA) for 30 min. The immunostaining was developed using 3,3-Diaminobenzidine (DAB) as chromogen and sections were counterstained with Mayer’s hematoxylin prior to imaging.

### Statistical analysis

Results are expressed as the mean ± 1 SD. A Shapiro-Wilk test for normality was performed on all datasets. Homogeneity was confirmed by a comparison of variances test. Parametric data was analyzed using either a Student’s t test for a two-group comparison, or a one-way analysis of variance when more than two groups were compared, followed by a post hoc Tukey test (GraphPad Software 9.0). \**P*<0.05 and \*\**P*<0.01 were considered significant. For *in vivo* 143B implantation studies, the sample size was calculated based on an anticipated effect size of 2.0 based on our *in vitro* studies comparing *CNTNAP4* KO and vector control. For this scenario, eight replicates per group was calculated to provide 80% power to detect effect sizes of at least 1.5, assuming a two-sided 0.05 level of significance.

### Study approval

All animals were housed, and procedures performed under a protocol approved by the IACUC of Johns Hopkins University (protocol MO21M112). Human samples were used under a written informed consent for tissue banking and Institutional Review Board (IRB) approval (protocol number IRB00119905).

## Data availability

The data generated in this study are available within the article and its supplementary data files. RNA sequencing data is freely available within the NCBI GEO database (GSE210373).

## Code availability

Our scRNA-seq analysis pipelines and codes can be obtained from the original author’s open source R package and script on Bioconducter (https://satijalab.org/seurat/). The default standard parameters are used in the analysis process.

## Acknowledgements

A.W.J. was funded by NIH/NIAMS (R01 AR070773, R01 AR079171, R01 DE031028), NIH/NIDCR (R21 DE027922), USAMRAA through the Peer-Reviewed Cancer Research Program (W81XWH-20-1-0302), Peer-Reviewed Medical Research Program (W81XWH-18-1-0121, W81XWH-18-1-0336), Peer-Reviewed Orthopaedic Research Program (W81XWH-20-1-0795) and Broad Agency Announcement (W81XWH-18-10613), American Cancer Society (Research Scholar Grant, RSG-18-027-01-CSM), and the Maryland Stem Cell Research Foundation. The content is solely the responsibility of the authors and does not necessarily represent the official views of the National Institute of Health or Department of Defense. We thank the JHU microscopy core facility, JHMI deep sequencing and microarray core facility, division of molecular pathology and Hao Zhang within the JHU Bloomberg Flow Cytometry and Immunology Core.

## Author contributions

Q.Q and S.R are co-first authors.

Conception and design: Q.Q., A.W.J.; acquisition, analysis, and interpretation of data: Q.Q., S.R., M.G.S., L.Z., M.C. and A.P.; donation of clinical samples: C.D.M., and E.F.M.; manuscript preparation: Q.Q., and S.R.; review and editing: X.Z. and A.W.J.; funding and final manuscript approval: A.W.J. All authors gave final approval of the completed version.

## Supplementary materials

### Supplementary Figures

**Supplementary Figure 1.**
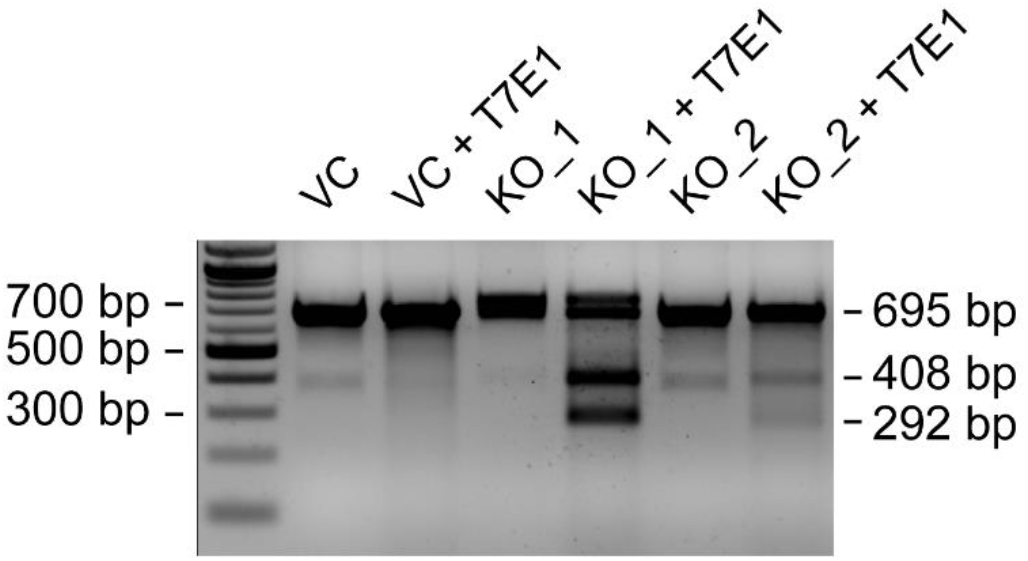
T7 endonuclease I assay in two *CNTNAP4* knockout (KO) single cell clones. The cleaved PCR products correspond to 408 and 292 bp, respectively.

**Supplementary Figure 2.**
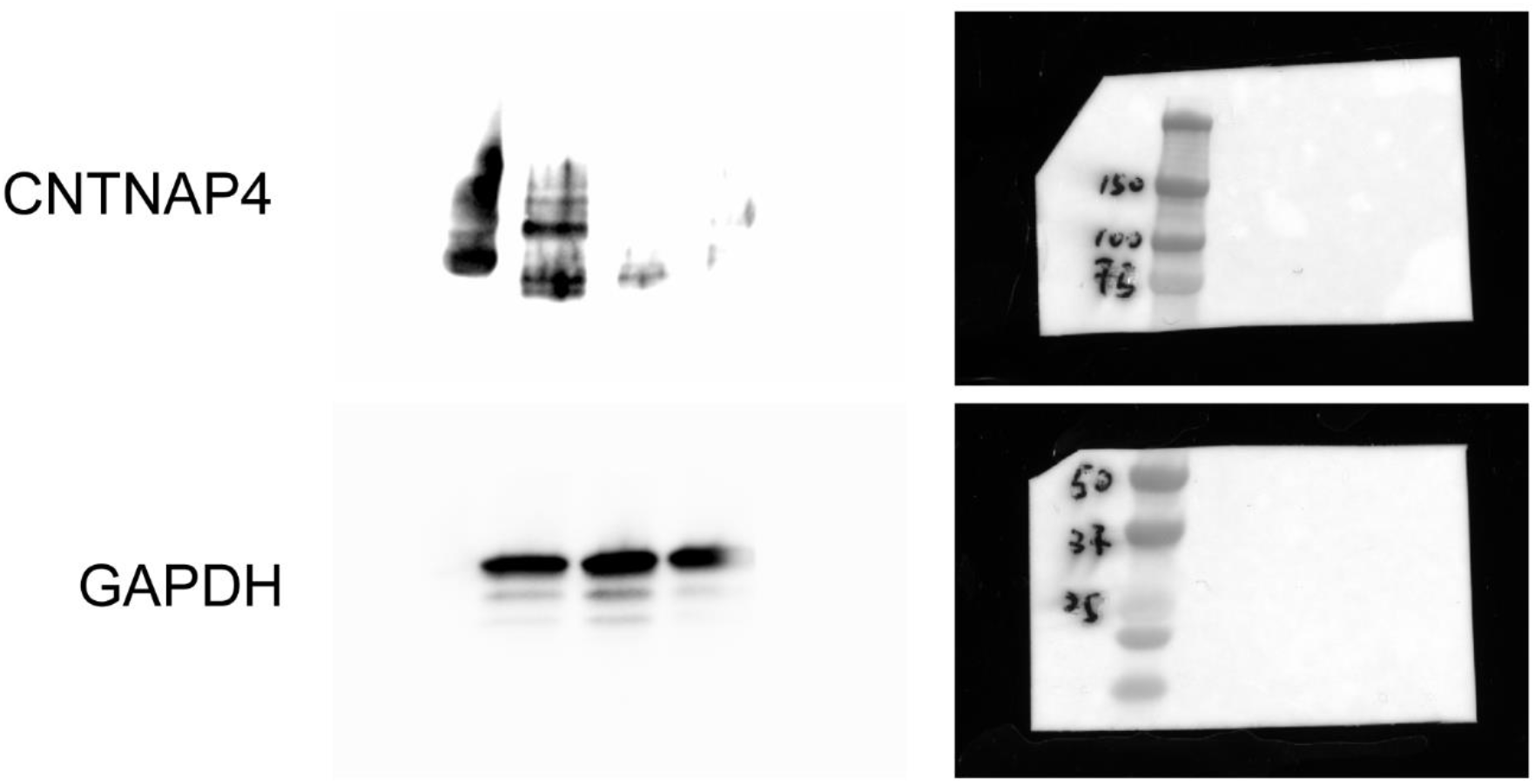
Full-size blots of Figure 1c.

**Supplementary Figure 3.**
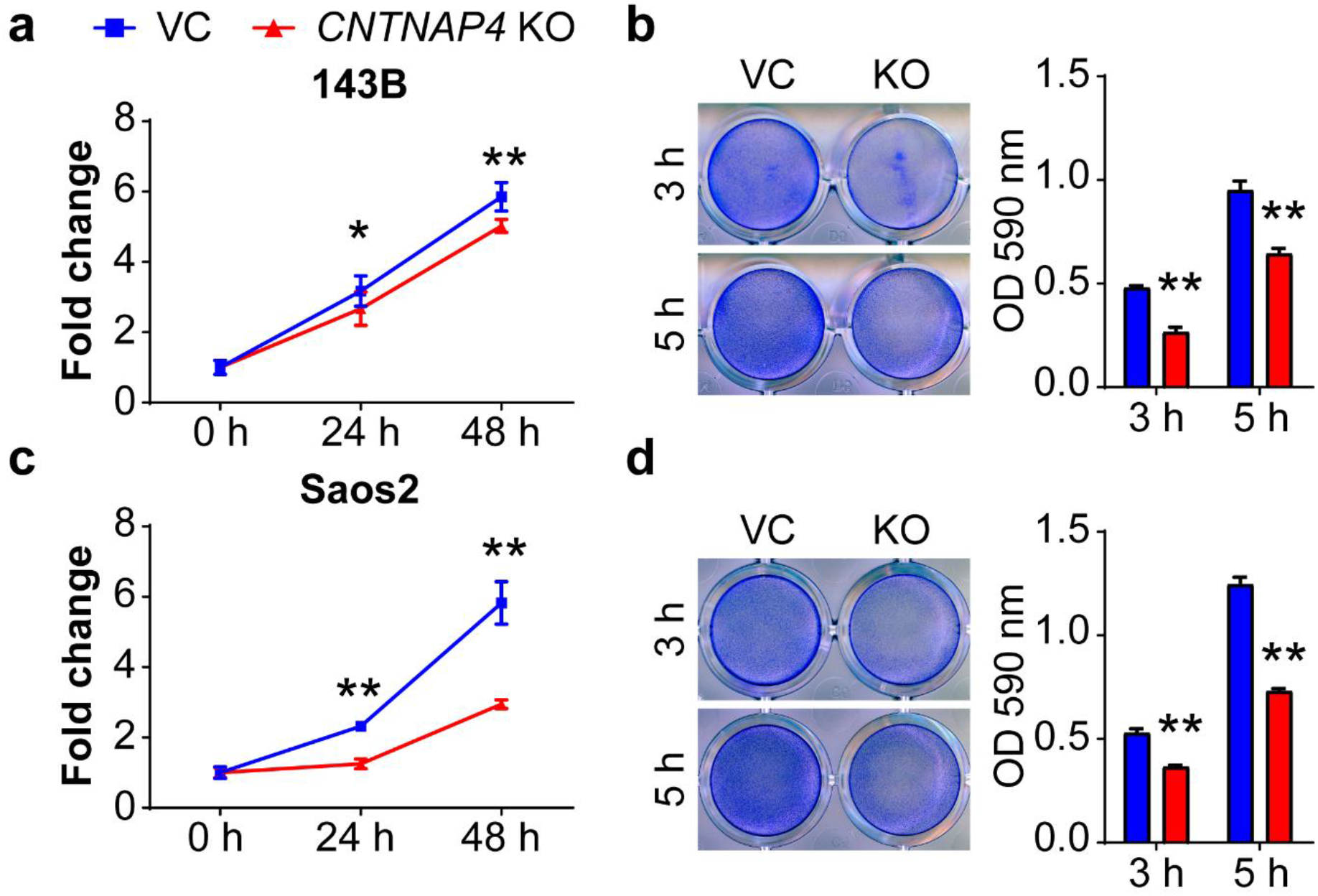
CRISPR-Cas9 mediated *CNTNAP4* KO in polyclonal 143B and Saos2 cells. **(a-b)** Effects of CRISPR-Cas9 mediated *CNTNAP4* gene deletion in polyclonal 143B OS cells. (**a**) Proliferation (MTS) assay with or without *CNTNAP4* KO (0-48 h) (**b**) Attachment assay as assessed by crystal violet staining (left) and quantification (right) with or without *CNTNAP4* KO (3-5 h). (**c-d**) Effects of CRISPR-mediated *CNTNAP4* gene deletion in polyclonal Saos2 OS cells. (**c**) Proliferation assay as assessed by MTS with or without *CNTNAP4* KO (0-48 h). (**d**) Attachment assay as assessed by crystal violet staining (left) and quantification (right) with or without *CNTNAP4* KO (3-5 h). Data shown as mean ± 1 SD. All experiments performed in triplicates, with results from a single replicate shown. \**P*<0.05, \*\**P*<0.01.

**Supplementary Figure 4.**
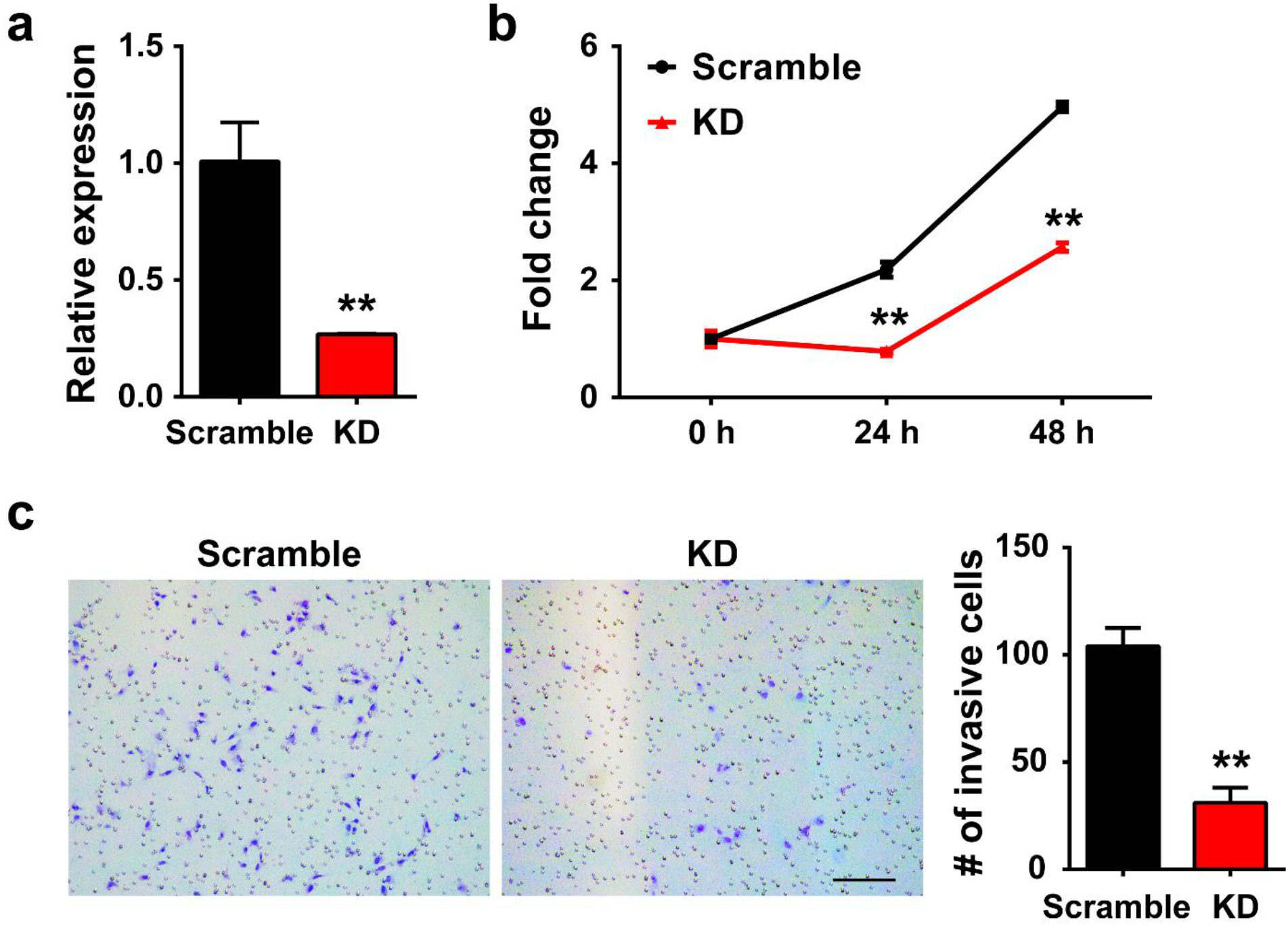
SiRNA mediated *CNTNAP4* KD in 143B cells. Effects of SiRNA mediated *CNTNAP4* gene knockdown in 143B OS cells in comparison to scramble control. (**a**) Confirmation of *CNTNAP4* knockdown efficiency by qPCR, 48 h after siRNA treatment. (**b**) Proliferation (MTS) with or without *CNTNAP4* KD (0-48 h). (**c**) Transwell invasion assay with crystal violet staining, with or without *CNTNAP4* KD (22 h). Data shown as mean ± 1 SD. All experiments performed in triplicates, with results from a single replicate shown. \*\**P*<0.01. Scale bar: 100 µm.

**Supplementary Figure 5.**
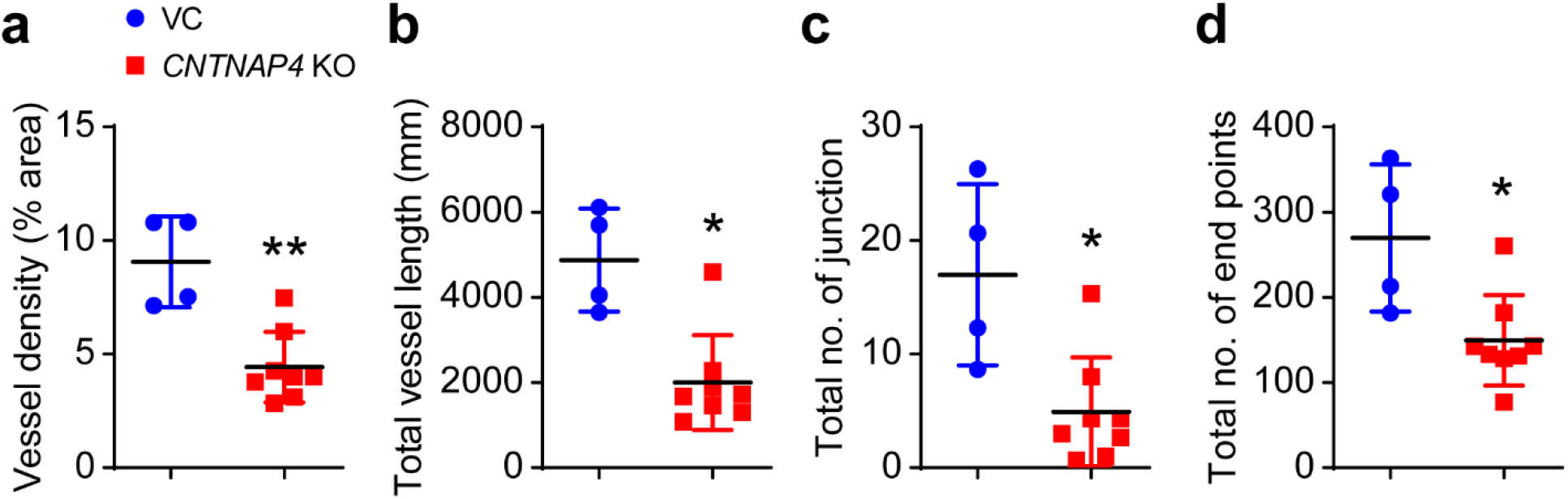
Vascular histomorphometric analysis in *CNTNAP4* VC and KO xenograft explants. (**a**) Vessel density (**b**) total vessel length (**c**) total number of junctions, and (**d**) total number of endpoints. VC (n=4) and *CNTNAP4* KO (n=8) tumor explants analyzed. Data shown as mean ± 1 SD, \*\**P*<0.01, \**P*<0.05.

**Supplementary Figure 6.**
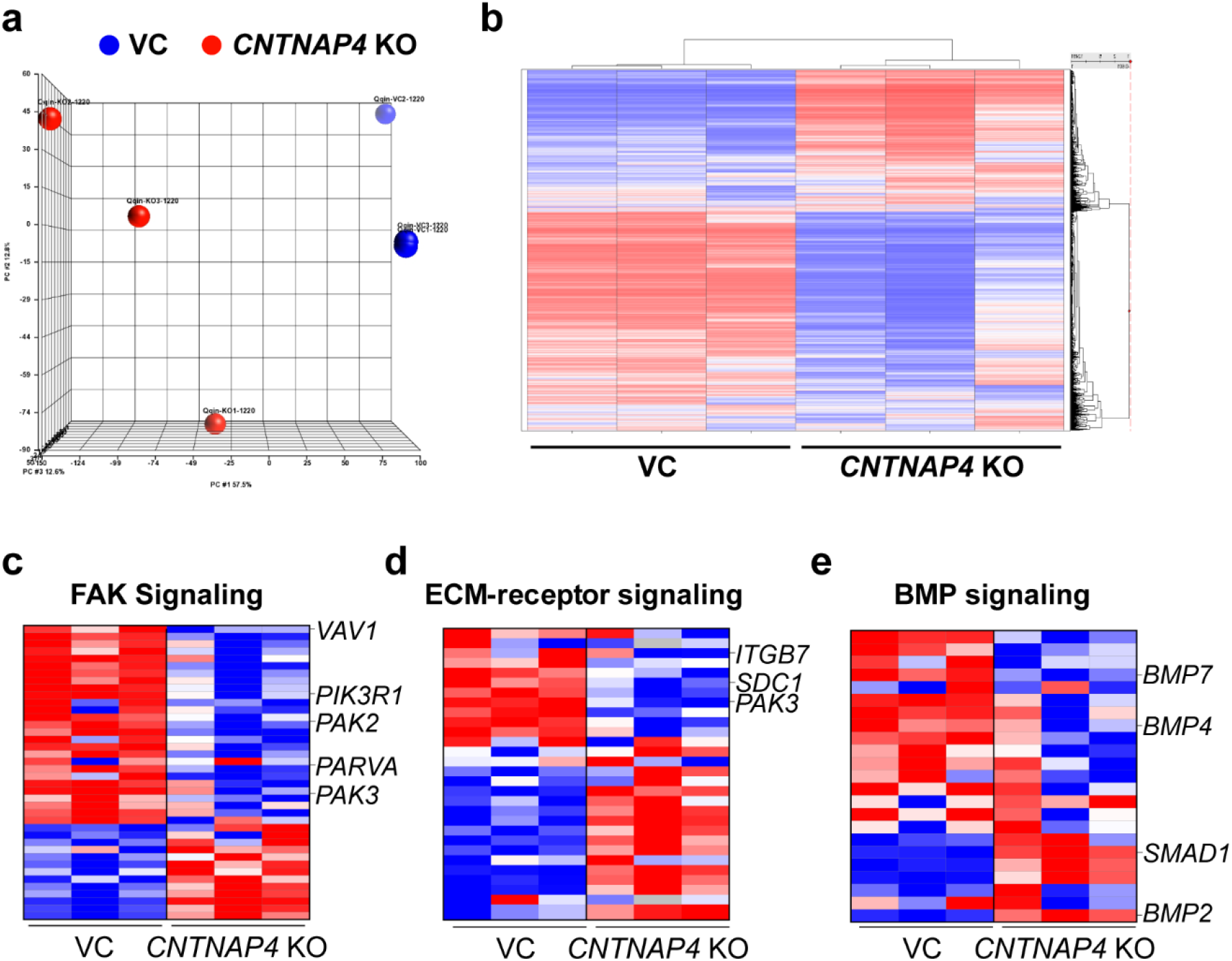
Additional transcriptomic analysis of clonal 143B cells with or without *CNTNAP4* gene deletion. (**a**) Principal component analysis among VC and *CNTNAP4* KO 143B osteosarcoma cells. (**b**) Clustering heatmap of all 19,565 protein coding genes expressed among VC and *CNTNAP4* KO 143B cells. (**c**) Heatmap of representative FAK signaling pathway related genes. Note that *CNTNAP4* KO decreased expression level of certain key genes (*VAV1, PIK3R1, PAK2, PARVA, PAK3)* with implications in cancer progression. (**d**) Heatmap of representative ECM-receptor signaling. Note that *CNTNAP4* KO decreased expression level of certain key genes (*ITGB7, SDC1, PAK3*) with implications in cancer progression. (**e**) Heatmap of representative BMP signaling.

**Supplementary Figure 7.**
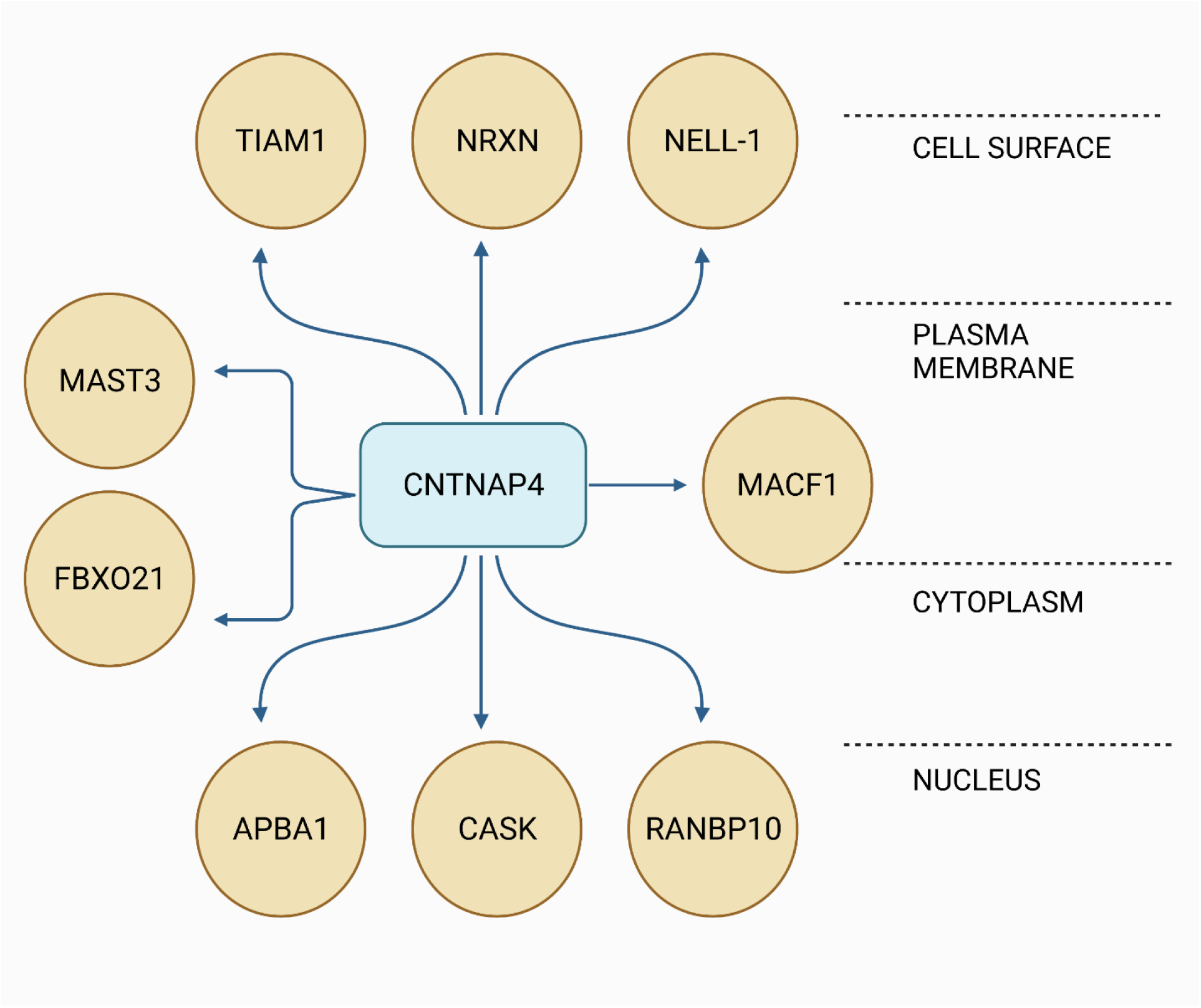
Protein-protein interaction of CNTNAP4 network obtained using the STRING and Innate DB database. TIAM1, NELL-1, MACF1, MAST3, FBX021, NRXN, APBA1, CASK, and RANBP10 physically interact with CNTNAP4. Cartoon created with BioRender.com.

**Supplementary Figure 8.**
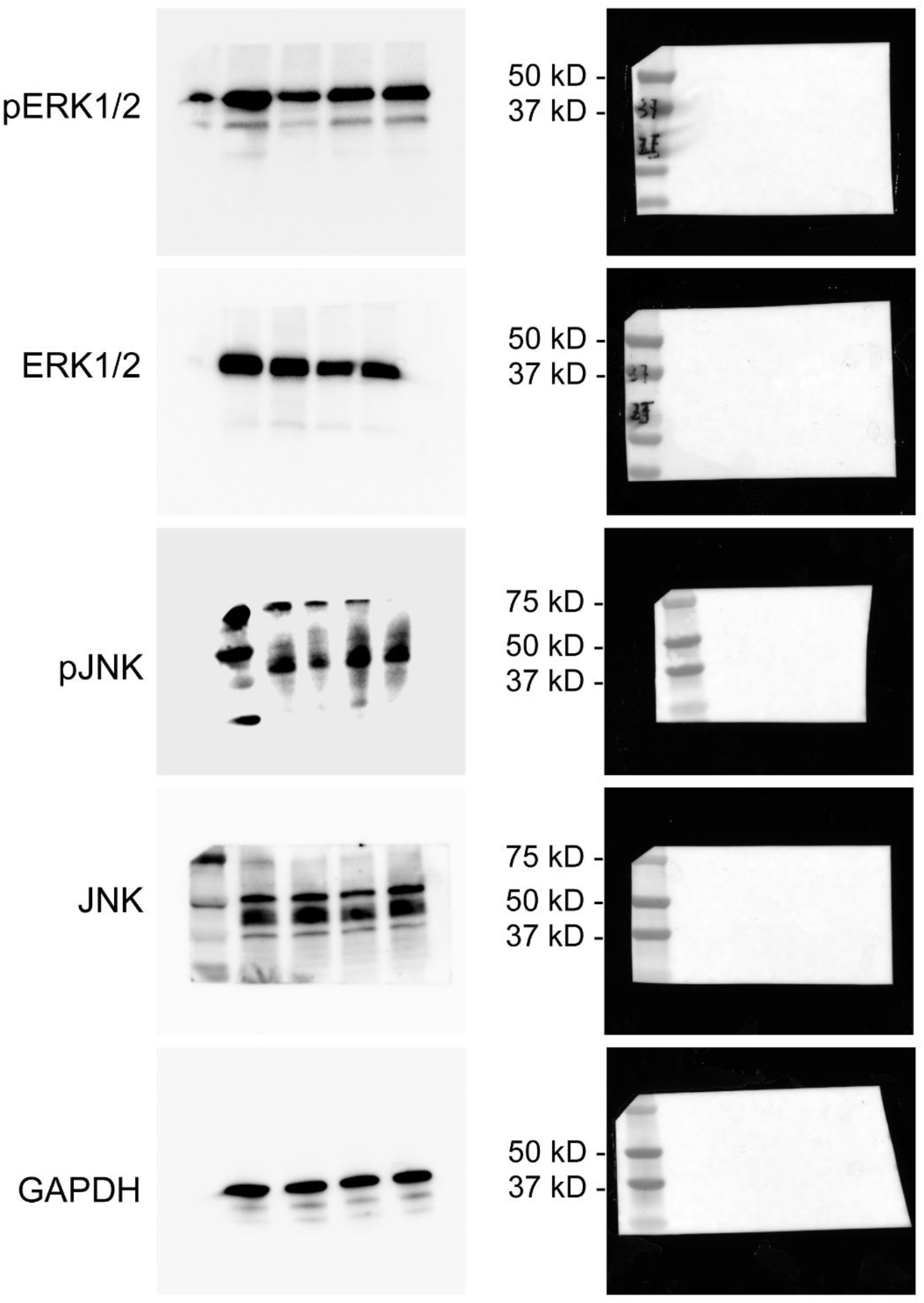
Full-size blots of Figure 4g.

**Supplementary Figure 9.**
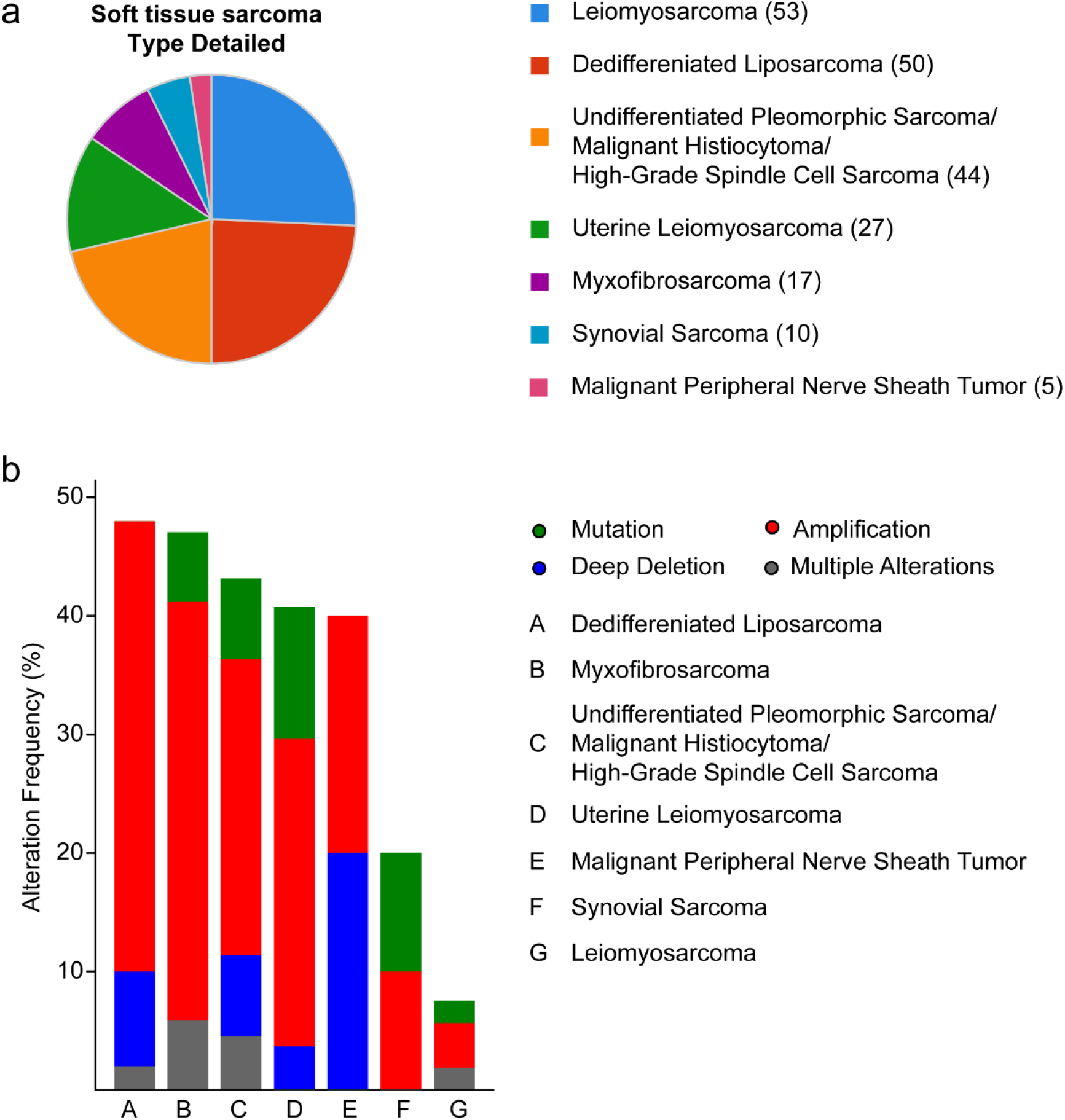
Cancer subtypes and frequency of all mutations in *CNTNAP4* associated genes. (**a**) Pie chart shows percentage of patients in each soft tissue sarcoma subtype (n=206). (**b**) Bar graph represents the distribution of alteration types of *CNTNAP4* associated genes in each soft tissue sarcoma subtype, including amplification (red), deep deletion (blue), multiple alterations (grey), and missense mutation (green).

**Supplementary Table 1.** Enriched Gene Ontology (GO) terms related to cancer signaling.

| **Downregulated GO terms_CNTNAP4 KO Vs VC** |  |  |
| --- | --- | --- |
| **GO term (Biological Processes)** | **Count** | ***p*-value** |
| GO:0000165~MAPK cascade | 90 | 0.004424 |
| GO:1901796~regulation of signal transduction by p53 class mediator | 65 | 1.73E-07 |
| GO:0051056~regulation of small GTPase mediated signal transduction | 61 | 1.30E-05 |
| GO:0090263~positive regulation of canonical Wnt signaling pathway | 53 | 0.001516 |
| GO:0030036~actin cytoskeleton organization | 53 | 0.031373 |
| GO:0048013~ephrin receptor signaling pathway | 36 | 1.86E-04 |
| GO:0007409~axonogenesis | 30 | 0.007502 |
| GO:0008543~fibroblast growth factor receptor signaling pathway | 27 | 0.04408 |
| GO:0006977~DNA damage response, signal transduction by p53 class mediator resulting in cell cycle arrest | 27 | 1.57E-04 |
| GO:0048010~vascular endothelial growth factor receptor signaling pathway | 23 | 0.031535 |
| GO:0030514~negative regulation of BMP signaling pathway | 23 | 0.037169 |
| GO:0034446~substrate adhesion-dependent cell spreading | 21 | 0.018607 |
| GO:1904263~positive regulation of TORC1 signaling | 10 | 0.031891 |
| GO:0046328~regulation of JNK cascade | 9 | 0.033764 |
| GO:0035331~negative regulation of hippo signaling | 9 | 0.002968 |
| GO:0060338~regulation of type I interferon-mediated signaling pathway | 7 | 0.006036 |
| GO:0034333~adherens junction assembly | 7 | 0.021732 |

**Supplementary Table 2.**
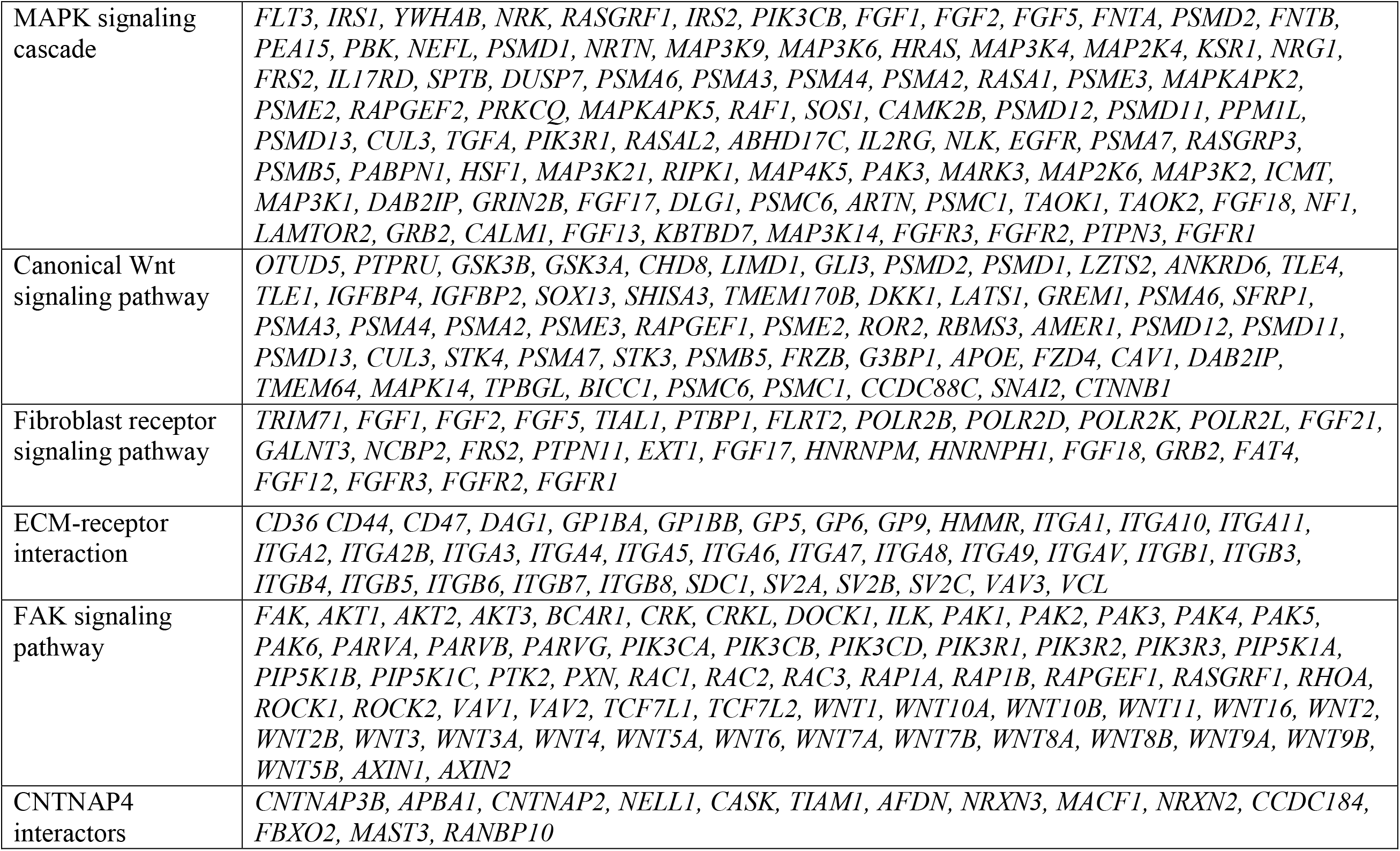
Gene list used for heatmap.

**Supplementary Table 3.**
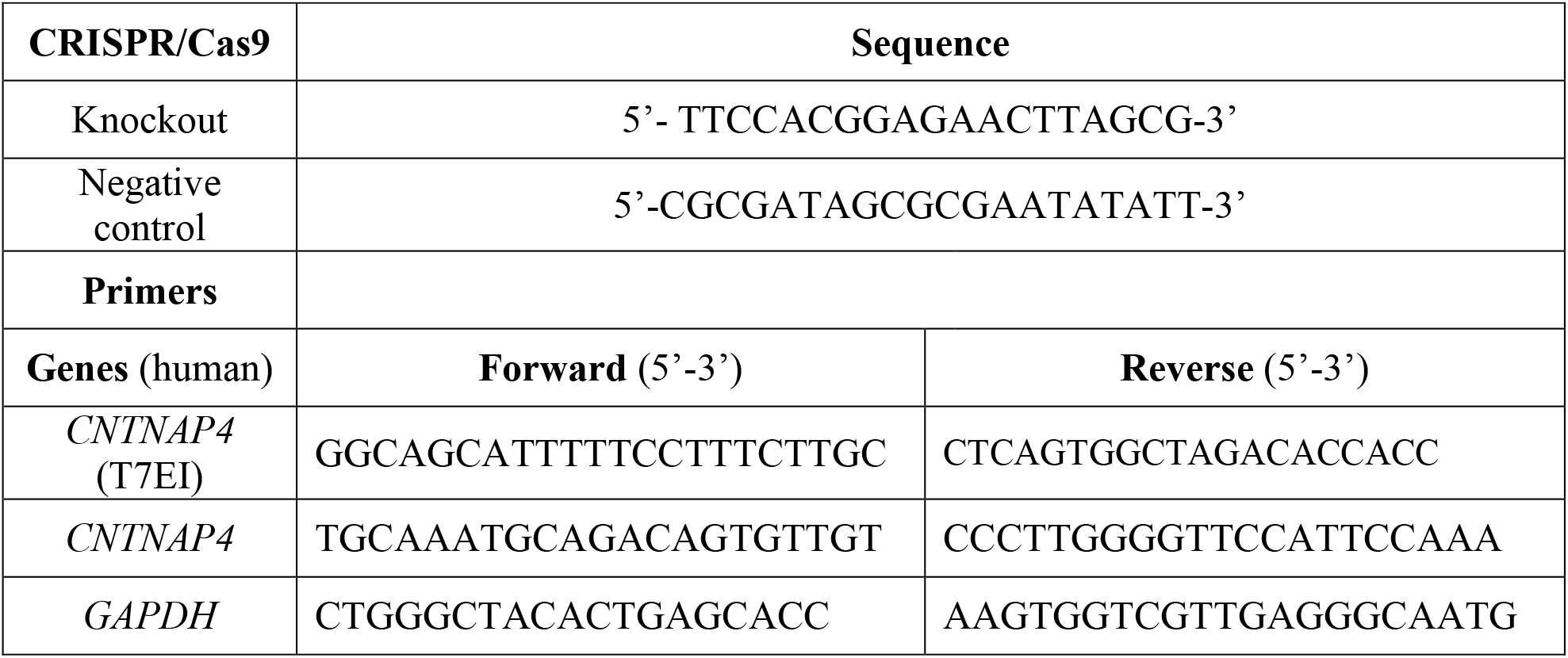
CRISPR/Cas9 sgRNA and primer sequences.

**Supplementary Table 4.** MAPK array protein targets layout.

|  | A | B | C | D | E | F | G | H |
| --- | --- | --- | --- | --- | --- | --- | --- | --- |
| 1 | POS | POS | NEG | NEG | Akt<br>(pS473) | CREB<br>(pS133) | ERK1<br>(pT202/Y204)<br>/ERK2<br>(Pt185/y187) | GSK3a<br>(pS21) |
| 2 |  |  |  |  |  |  |  |  |
| 3 | GSK3b<br>(pS9) | HSP27<br>(pS82) | JNK<br>(pT183) | MEK<br>(pS217/T221) | MKK3<br>(pS189) | MKK6<br>(pS207) | MSK2<br>(pS360) | mTOR<br>(pS2448) |
| 4 |  |  |  |  |  |  |  |  |
| 5 | p38<br>(pT180/Y182) | p53<br>(pS15) | P70S6K<br>(pT421/S424) | RSK1<br>(pS380) | RSK2<br>(pS386) | NEG | NEG | POS |
| 6 |  |  |  |  |  |  |  |  |

**Supplementary Table 5.** List of antibodies used.

| **Antibody** | **Company** | **Catalog #** | **Use** | **Dilution** |
| --- | --- | --- | --- | --- |
| Rabbit anti-CNTNAP4 | Biorbyt | orb544737 | IHC, WB | 1:100 |
| Mouse anti-Human Nuclei | Millipore-Sigma | MAB1281 | IF | 1:200 |
| Rabbit anti-CD31 | Abcam | ab28364 | IF | 1:200 |
| Rabbit anti-Ki67 | Abcam | ab15580 | IF | 1:200 |
| Goat anti-Mouse AF488 | Abcam | ab150117 | IF | 1:1000 |
| Goat anti-Rabbit AF488 | Abcam | ab150077 | IF | 1:1000 |
| Goat anti-Rabbit DyLight 594 | Vector Laboratories | DI-1594 | IF | 1:1000 |
| Goat anti-Rabbit, HRP | Invitrogen | 32460 | IHC | 1:1000 |
| Rabbit anti-p44/42 MAPK (Erk1/2) | Cell Signaling Technology | 9102 | WB, IF | 1:100 |
| Rabbit anti-Phospho-p44/42 MAPK (Erk1/2) | Cell Signaling Technology | 9101 | WB, IF | 1:100 |
| Rabbit anti-FGF | Cell Signaling Technology | 61997 | WB | 1:100 |
| Rabbit anti-JNK | Cell Signaling Technology | 9252 | WB | 1:100 |
| Mouse anti- Phospho-JNK | Cell Signaling Technology | 9255 | WB | 1:100 |
| Rabbit anti-GAPDH | Cell Signaling Technology | 5174 | WB | 1:100 |
| Anti-mouse IgG, HRP-linked Antibody | Cell Signaling Technology | 7076 | WB | 1:2500 |
| Anti-biotin, HRP-linked Antibody | Cell Signaling Technology | 7075 | WB | 1:2500 |
| Anti-rabbit IgG, HRP-linked Antibody | Cell Signaling Technology | 7074 | WB | 1:2500 |
| IF: Immunofluorescent staining; IHC: Immunohistochemistry; WB: Western Blot |  |  |  |  |

